# The hepatocyte Epidermal Growth Factor Receptor (EGFR) pathway regulates the cellular interactome within the liver fibrotic niche

**DOI:** 10.1101/2023.11.03.565317

**Authors:** Ester Gonzalez-Sanchez, Javier Vaquero, Daniel Caballero-Diaz, Jan Grzelak, Noel P Fusté, Esther Bertran, Josep Amengual, Juan Garcia-Saez, Beatriz Martín-Mur, Marta Gut, Anna Esteve-Codina, Ania Alay, Cedric Coulouarn, Silvia Calero, Pilar Valdecantos, Angela M. Valverde, Aránzazu Sánchez, Blanca Herrera, Isabel Fabregat

## Abstract

**Background & Aims:** Liver fibrosis is the consequence of chronic liver injury in the presence of an inflammatory component. Although the main executors of this activation are known, the mechanisms that lead to the inflammatory process that mediates the production of profibrotic factors are not well characterized. The Epidermal Growth Factor Receptor (EGFR) signaling in hepatocytes is essential for the regenerative process of the liver; however, its potential role in regulating the fibrotic niche is not yet clear.

**Approach & Results:** Our group generated a mouse model that expresses an inactive truncated form of the EGFR specifically in hepatocytes (ΔEGFR mice). Here, we have analyzed the response of WT and ΔEGFR mice to chronic treatment with CCl_4_.

**Results:** indicated that the hallmarks of liver fibrosis were attenuated in CCl_4_-treated ΔEGFR mice when compared to WT mice, coinciding with a faster resolution of the fibrotic process and an ameliorated damage. The absence of EGFR activity in hepatocytes induced changes in the pattern of immune cells in the liver, with a notable change in the population of M2 macrophages, more related to fibrosis resolution, as well as an increase in the population of lymphocytes related to eradication of the damage. Transcriptomic analysis of hepatocytes and secretome studies from extracellular media in *in vitro* studies allowed to elucidate the specific molecular mechanisms regulated by EGFR that mediate hepatocyte production of both pro-inflammatory and pro-fibrotic mediators.

**Conclusions:** Our results support a pro-inflammatory and pro-fibrogenic role for the hepatocyte EGFR pathway during chronic liver damage.

## INTRODUCTION

Fibrosis is a highly conserved and coordinated, initially protective, response to tissue injury. The interaction among multiple pathways determines whether fibrosis is self-limiting and homeostatic, or whether it is uncontrolled and excessive.(1) When a chronic injury takes place in the liver, mobilization of lymphocytes and other inflammatory cells occurs, thus setting the stage for persistence of an inflammatory response. Macrophages produce profibrotic mediators that are responsible for the activation of quiescent HSC to a myofibroblast (MFB) phenotype. MFBs are the principal source of extracellular matrix (ECM) protein accumulation and prominent mediators of fibrogenesis.(2)

The Epidermal Growth Factor Receptor (EGFR) or ErbB-1 is a classical tyrosine kinase receptor that belongs to the family of ErbB receptors and is highly expressed in the liver. The EGFR pathway plays essential roles in hepatocyte proliferation, which is highlighted from the fact that, together with the Hepatocyte Growth Factor (HGF), EGFR ligands are the only mitogens for hepatocytes, playing important roles in liver regeneration and hepatocellular carcinogenesis.(3) However, this is not the only role that this pathway plays in the liver. In fact, the EGFR signaling system has been identified as a key player in all stages of the liver response to injury, from early inflammation and hepatocellular proliferation to fibrogenesis and neoplastic transformation.(4) The role of EGFR in chronic liver injury and fibrosis is also relevant for human pathophysiology, since EGF, EGFR and phospho-EGFR levels increase in liver of cirrhotic or NAFLD (non-alcoholic fatty liver disease) patients.(3) A direct role of EGFR, or its ligands, in hepatic stellate cell (HSC) activation and subsequent extracellular matrix modification has been suggested, concomitant with regulation of steatosis, in different *in vitro* and *in vivo* animal models of liver fibrosis and NAFLD.(5–9) However, different studies pointed out that the EGFR signaling might be protective in other processes of liver injury and fibrosis, due to its participation in the physiological regulation of bile acid synthesis.(10, 11)

With the aim of better understanding the specific role of the hepatocyte EGFR pathway in liver physiology and pathology, some years ago we generated a transgenic mouse model expressing a hepatocyte-specific truncated form of human EGFR, which acts as negative dominant mutant (ΔEGFR) and allows definition of its tyrosine kinase-dependent functions.(12) One of the most interesting results found in this mouse model was that EGFR catalytic activity is critical in the early preneoplastic stages of the liver because ΔEGFR mice showed a delay in the appearance of diethyl-nitrosamine (DEN)-induced tumors, which correlated with a strong attenuation and delay in the DEN-induced inflammatory process.(12) Previous studies had suggested that pro-inflammatory stimuli could mediate transactivation of the EGFR pathway, which would contribute to carcinogenesis in inflammatory liver diseases.(13, 14) However, our data pointed out to a role of the EGFR pathway in hepatocytes upstream of the inflammatory process. In this context, further investigations were required to comprehensively study the direct role of EGFR in hepatocytes in regulating inflammation during chronic liver injury.

Accordingly, the aim of this work was to elucidate whether the catalytic activity of the EGFR in hepatocytes could participate in regulating the inflammatory/pro-fibrotic environment in response to a chronic liver insult.

## METHODS

### In vivo experimental procedures

A novel transgenic mouse model that expresses a hepatocyte-specific truncated form of human EGFR (***Suppl. Fig. 1A***), which acts as dominant negative mutant (ΔEGFR) and allows definition of its tyrosine kinase-dependent functions, was used.(2) Chronic liver injury was induced by carbon tetrachloride (CCL_4_) treatment. For this, 0.48 g/kg body weight CCl_4_ (1:10 v/v in mineral oil), or vehicle, were administered in 8 week-old C57BL/6 male mice by intraperitoneal injection (3 µl/g body weight) twice a week for 4 or 8 weeks. At these timepoints, animals were sacrificed, and blood and liver tissue were collected (***Suppl. Fig. 1B***). Transgenic ΔEGFR mouse line is maintained in heterozygosity, therefore WT and ΔEGFR mice belong to the same strain. WT and ΔEGFR mice used for experiments belonged to the same littermates. We proved that ΔEGFR mice continued expressing the human truncated transgene after the CCl_4_ treatment (***Suppl. Fig. 1C***). Wild Type (WT) and ΔEGFR mouse lines were maintained in a C57BL/6 background in the UCM animal facility allowed food and water *ad libitum* in temperature-controlled rooms under a 12h light/dark cycle, and routinely screened for pathogens in accordance with Federation of European Laboratory Animal Science Associations procedures. All animal procedures were done conformed to ARRIVE Guidelines and European Union Directive 86/609/EEC and Recommendation 2007/526/ EC, enforced in Spanish law under RD 1201/2005. Animal protocols including sample size decisions and randomization and blinding strategies were approved by the Animal Experimentation Ethics Committee of the UCM and the Animal Welfare Division of the Environmental Affairs Council of the Government of Madrid (Proex 262.6/21). We have followed the 3R recommendations: Replacement, Reduction and Refinement. To avoid potential bias, experimental groups included animals from different breedings. Furthermore, littermates included mice treated with mineral oil and CCl4.

### Histological and immunohistochemical analyses

Liver sections (3-10 μm) were cut from paraffin. Hematoxylin-eosin staining and immunohistochemical analyses were performed as previously described.(12) Liver sections were stained with Sirius red to evaluate collagen deposition and therefore fibrosis. Primary antibodies (listed in ***Suppl. Table 1***) were incubated overnight at 4°C and binding developed with the Vectastain ABC kit (Vector Laboratories, Burlingame, CA, USA). Nuclei were stained with hematoxylin solution and preparations were mounted in DPX. Stained sections were visualized in a Nikon Eclipse 80i microscope coupled to a Nikon DS-Ri1 digital camera or scanned on a virtual slide scanner NanoZoomer 2.0 HT (Hamamatsu, Tokyo, Japan) at the Histopathology Facility of Institute for Research in Biomedicine -IRB (Barcelona, Spain). Morphometric analyses were performed blinded, using ImageJ analysis software (National Institutes of Health, Bethesda, MD, USA).

In parallel, serum levels of the hepatocellular damage markers aspartate aminotransferase (AST) and alanine aminotransferase (ALT) were measured in Echevarne Laboratories (Barcelona, Spain).

### Isolation of hepatic non-parenchymal cells for immune cell populations analysis

Livers were collected and washed with PBS. Immediately, they were transferred to HBSS (Hank’s Balanced Salt solution; Gibco, Life Technologies, Carlsbad, CA, USA) at RT, disintegrated and filtered through 100 μm cell strainers. Then, homogenates were centrifuged at 500 x g for 5 min at RT and cell pellets were resuspended in a 36% Percoll solution (GE Healthcare Bio-Sciences AB, Uppsala, Sweden) containing 100 UI/mL of heparin (HIBOR 5000 UI, ROVI, Madrid, Spain). After centrifugation at 800 x g without brake for 20 min at RT, supernatants were discarded and then erythrocytes were removed from cell pellets by using a red blood cell lysis buffer (150 mM NH_4_Cl, 10 mM KHCO_3_, 0.1 mM EDTA pH 7.3). The resulting cell pellets were washed with cold HBSS, centrifuged at 500 x g for 5 min at 4° C and, finally, cells were resuspended in cold HBSS for further analysis. Isolated non-parenchymal liver cells were incubated with the antibodies listed in ***Suppl. Table 2***, or their corresponding isotype controls for 20 min at RT protected from light. After washing steps, cells were resuspended with PBS. Flow cytometry data were acquired with a FACSCanto II and data analysis was performed using Cytomics FC500 with the CXP program. One-way ANOVA to calculate p-values once normal distribution of data was verified using Shapiro-Wilk test.

### Analysis of gene expression by RT-qPCR

mRNA expression levels of cell type markers, pro-/anti-inflammatory factors and genes involved in collagen homeostasis were analyzed by Real time quantitative PCR (RT-qPCR). Total RNA was isolated from the different cells or tissues using “RNeasy Mini Kit” (Qiagen, Hilden, Germany). cDNA was produced using the High-capacity cDNA Reverse Transcription Kit (Applied Biosystems, Waltham, MA, USA). RT-qPCR was performed in duplicate in an ABIPrism7700 System following manufacturer’s protocol. SYBR Green PCR Master Mix was used for PCR reactions (Applied Biosystems). Primers are listed in ***Suppl. Table 3*** and ***4*.**

### RNA Sequencing (RNA-seq) analysis in hepatocytes

Primary mouse hepatocytes were isolated as previously described.(15) Briefly, livers from 2-3 months old male mice were perfused with Hank’s balanced salt solution supplemented with 10 mM Hepes and 0.2 mM EGTA for 5 min, followed by a 15 min perfusion with William’s medium E containing 10 mM Hepes and 0.03% collagenase type 1 (125 U/mg; Worthington). Livers were further minced, filtered through a 70 μm cell strainer (BD Biosciences Franklin Lakes, NJ, USA) and viable hepatocytes were selected by centrifugation in Percoll and stored at -80°C. Total RNA from *Mus musculus* was quantified by Qubit® RNA BR Assay kit (Thermo Fisher Scientific, Bremen, Germany) and the RNA integrity was estimated by using RNA 6000 Nano Bioanalyzer 2100 Assay (Agilent, Santa Clara, CA, USA). The RNA-seq libraries were prepared with KAPA Stranded mRNA-Seq Illumina® Platforms Kit (Roche) following the manufactureŕs recommendations starting with 500 ng of total RNA as the input material. The library was quality controlled on an Agilent 2100 Bioanalyzer with the DNA 7500 assay. The libraries were sequenced on NovaSeq 6000 (Illumina) with a read length of 2x151bp, following the manufacturer’s protocol for dual indexing. Image analysis, base calling and quality scoring of the run were processed using the manufacturer’s software Real Time Analysis (RTA 3.4.4).

RNA-seq reads were mapped against *Mus musculus* reference genome (GRCm39) using STAR aligner version 2.7.8a(16) with ENCODE parameters. Annotated genes were quantified with RSEM version 1.3.0(17) with default parameters, using the annotation file from GENCODE version M31. Differential expression analysis was performed with limma v3.4.2 R package, using TMM normalization. The voom function(18) was used to estimate mean-variance relationship and to compute observation-level weights. The linear model was fitted with the voom-transformed counts and contrasts were extracted. Genes were considered differentially expressed (DEG) with a p-value adjusted < 0.05 and subsets of DEG were represented in heatmaps with the pheatmap R package, using voom-transformed counts scaled by row. A functional enrichment analysis was performed on the DEG with gprofiler2 v0.1.8(19) using ENSEMBL databases as reference. Additionally, a gene set enrichment analysis (GSEA) was performed with the list of pre-ranked genes by a t-statistic with the R package fgsea v1.12.0,(20) against mouse Reactome database and M5 gene set collection from MSigDB.

### Western blot analysis

Western blot was performed as previously described.(21) Briefly, tissue and cells were lysed in RIPA lysis buffer supplemented with orthovanadate and a cocktail of protease inhibitors at 4°C (using a TissueLyser (Qiagen) to homogenize liver tissue) and protein concentrations determined using BCA kit (Pierce). Proteins were separated by SDS electrophoresis on 10% polyacrylamide gels and transferred to nitrocellulose membranes. Membranes were incubated with the primary antibodies (***Suppl. Table 1***) overnight at 4°C and then with ECL Mouse IgG and Rabbit IgG, HRP-Linked antibodies (GE Healthcare, Buckinghamshire, UK) (1/2000) for 1 h at room temperature. Blots were visualized using ChemiDoc™ Touch Imaging System (BioRad, Hercules, CA, USA) and densitometric analysis was performed using Image Lab™ Software (BioRad).

### In vitro analyses

Immortalized hepatocytes were isolated from ΔEGFR or WT mice as previously described.(12) Hep3B human hepatocarcinoma cells and THP-1 human monocytes were obtained from the European Collection of Cell Cultures whereas RAW 264.7 mouse macrophage cell line was kindly provided by Dr. Nicolas Chignard (CRSA, Paris, France). Cells were cultured with DMEM medium supplemented with 10% (v/v) heat inactivated FBS, 100 U/mL penicillin and 100 μg/mL streptomycin, maintained in a humidified atmosphere of 37°C, 5% CO_2_ and routinely screened for the presence of mycoplasma. Conditioned media were prepared as follows: murine immortalized hepatocytes or Hep3B cells were seeded and 24 h later serum starved overnight. The cells were then treated with heparin-binding EGF like growth factor (HB-EGF, an EGFR ligand) at a concentration of 20 ng/ml. PBS was used as vehicle. After 30 min, the cells from one plate of each cell line and experimental condition were collected for control studies to validate the activation of the EGFR pathway. Two hours after the treatment, the medium was replaced by FBS-free medium in the absence of the EGFR ligand. The conditioned media were collected 24 h later and used to analyze their effects on RAW 264.7 mouse macrophages or THP-1 human monocytes. Gene expression was determined by RT-qPCR in RAW 264.7 and THP-1 monocytes after 48 h-incubation with conditioned media collected from murine immortalized hepatocytes or Hep3B cells, respectively. In the case of THP-1 monocytes, the effect of conditioned media from Hep3B cells on cell adhesion was evaluated after 24 h of exposure with the xCELLigence System (Agilent,), using phorbol myristate acetate (PMA), as positive control.

### Proteomic analysis of the secretome

Proteomic analysis of the conditioned media obtained in the *in vitro* experiments explained above was performed in the Proteomics Unit of the Complutense University of Madrid. A label-free experiment was conducted. Briefly, secretome samples were concentrated with speed vac and resuspended in urea 8M. Proteins were digested using an iST kit (Preomics, Planegg, Germany). The resulting peptides were analyzed using liquid nano-chromatography (Vanquish Neo, Thermo Fisher Scientific), coupled to high-resolution mass spectrometer Q-Exactive HF (Thermo Fisher Scientific,). Proteins were identified using Proteome Discover 3.0 software (Thermo Scientific) and the search engine Mascot 2.6 (matrixscience.com). The database used was Uniprot (UP-000000589). For quantitative proteomics, using the aforementioned software, chromatograms and retention times of all samples were aligned. Afterwards, total protein abundance between different samples is normalized. Statistically, the Student’s t-test is applied to assess proteins differential abundance between samples. *p* values of less than 0.05 were considered statistically significant.

### Data analysis of publicly available human gene expression

Fujiwara et al. cohort of liver biopsies from HCC-naïve NAFLD patients was accessed through Gene Expression Omnibus (GEO) accession number GSE193066. Relative log-expression normalized data was directly downloaded from GEO. Two EGFR signaling gene signatures were obtained from MSigDB v2023.1:(22) “EGF/EGFR signaling” from WikiPathways (C2 CP) and “EGFR target genes” from TFT (C3) and gene set variation analysis (GSVA)(23) was used to assess the relative activation of the signature in the samples.

Correlation in fibrotic samples between the EGFR signature and genes of interest was assessed using Pearson correlation. Kendall’s τ was used to assess the association between either EGFR signatures or gene expression with fibrosis stage. All p-values were adjusted using Bonferroni correction test. All analyses were performed using R v4.0.4.(24)

### Statistical analysis

Statistical analyses have been specified in each of the technologies. Data representation as box-whisker plots were performed using GraphPad Prism software version 5.0 (GraphPad Software San Diego, CA, USA). Once a normal distribution of data was verified using Shapiro-Wilk test, differences between groups were compared using Student’s t-test or ANOVA and considered statistically significant when p < 0.05.

## RESULTS

### Inactivation of EGFR in hepatocytes ameliorates CCl_4_-induced liver damage and fibrosis

We first examined the consequences of inactivating EGFR signaling in hepatocytes on the liver damage progression after 4 or 8 weeks CCl_4_ chronic treatment. As observed in ***Fig. 1A***, serum levels of transaminases increased after 8 weeks of treatment in WT mice (no differences were observed at 4 weeks of treatment, results not shown). However, no significant increase was observed in ΔEGFR mice. Histological analysis also revealed less damage in the liver parenchyma after 8 weeks of treatment (***Fig. 1B***), correlating with decreased levels of Alpha-Smooth Muscle Actin (a-SMA), characteristic of activated MFB (***Fig. 1C-D***). In ΔEGFR livers the expression and deposits of a-SMA increased after 4 weeks but decreased at 8 weeks, reflecting resolution of fibrosis. Specific analysis of the process of fibrosis revealed that the increase in the deposits of collagen fibers, caused by the CCl_4_ treatment, was significantly lower in ΔEGFR livers (***Fig. 2A***). The expression of different collagen genes, such *Col1a1* or *Col3a1*, significantly increased after 4 weeks of treatment; however, increases in expression were lower in ΔEGFR livers, which was particularly relevant in *Col3a1* (***Fig. 2B***). Together with differences in collagen genes expression, it was very interesting to find differences in lysyl-oxidase related genes, responsible for the chain inter-polypeptide cross-link of collagens, which were significantly increased in WT but not in ΔEGFR livers after 4 weeks of CCl_4_ treatment (***Fig. 2B***). Also, worthy to mention that ΔEGFR mice presented different patterns in the expression of metalloproteases, particularly *Mmp2* and *Mmp9*, with a potential role in the resolution of fibrosis, which were significantly increased in ΔEGFR, but not in WT livers, after 8 weeks of treatment (***Fig. 2C***). Other metalloproteases analyzed (*Mmp12*, *Mmp13*) did not suffer any changes in WT or ΔEGFR livers (results not shown). The expression of *Timp1* (Metalloprotease Inhibitor 1) was significantly lower in ΔEGFR livers after 4 weeks of CCl_4_ treatment (***Fig. 2C***), which would indicate a higher metalloprotease activity. We could not find relevant differences among WT and ΔEGFR in the expression of genes related to the Transforming Growth Factor-beta (TGF-β), as one of the major pro-fibrotic factors. Thus, although the increase in some of its ligands appeared to be delayed in ΔEGFR mice (***Suppl. Fig. 2A***), phosphorylation of Smad3, as a hallmark of the activation of the pathway, was significantly increased after 8 weeks of treatment in both WT and ΔEGFR livers (***Suppl. Fig. 2B***).

**Figure 1.**
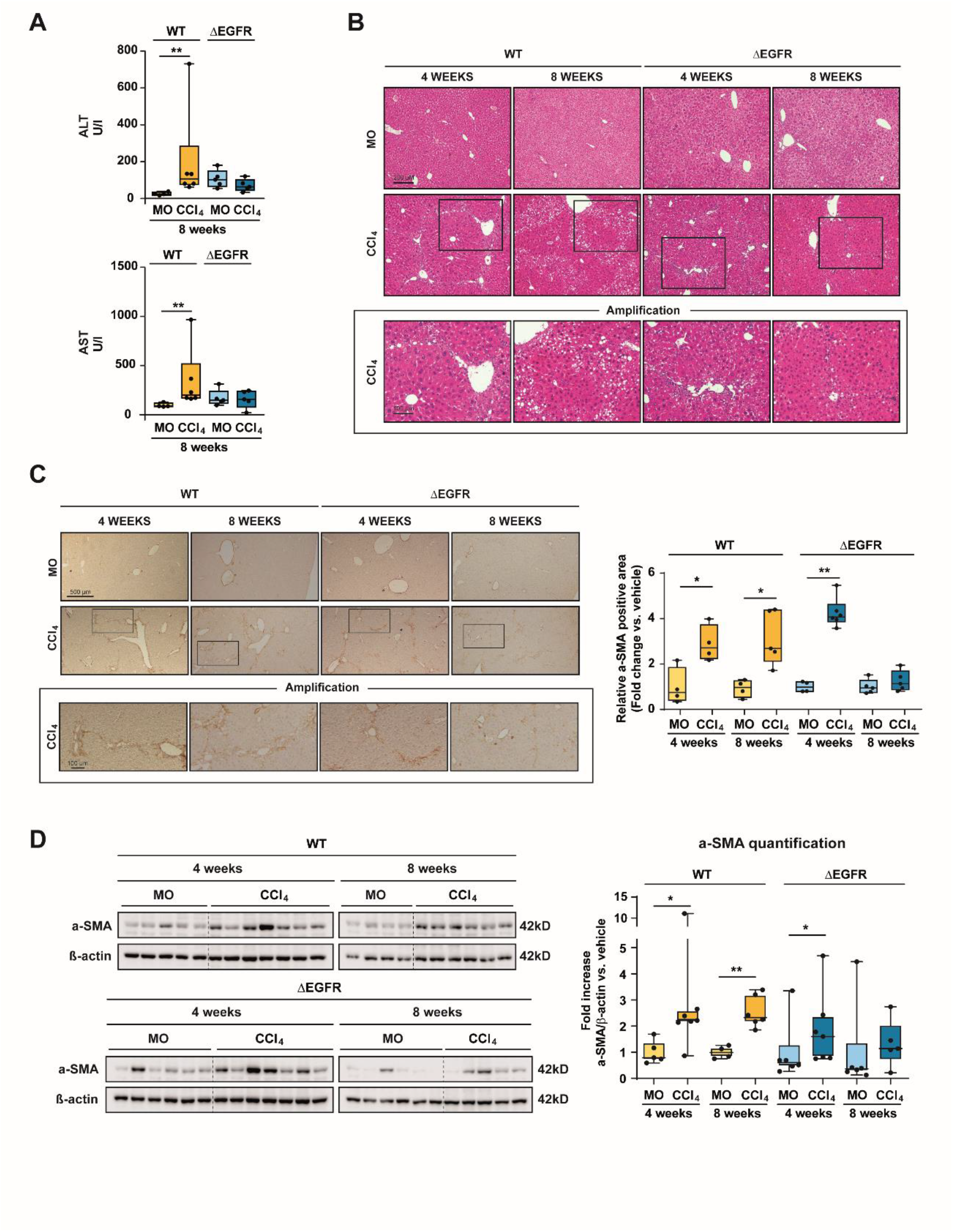
The impairment of EGFR catalytic activity in hepatocytes attenuates CCl_4_-induced liver damage and myofibroblast activation. A-B. Serum concentrations of alanine (ALT) and aspartate aminotransferase (ASP) (A) and hematoxylin and eosin-stained liver tissue sections (B) were analyzed to determine CCl_4_-induced liver damage (n=4-6 mice per group). C-D. The levels of Alpha-Smooth Muscle Actin (a-SMA), a marker of activated myofibroblasts, was evaluated by immunostaining (C) (quantification of the positive-stained area on the right) and western blot (D) (densitometric analysis respect to β-actin as loading control on the right). Data from 4-7 animals per group are presented as box-and-whisker plots. For panels C and D data are expressed as fold change versus vehicle (mineral oil: MO). *p<0.05, **p<0.01 using Student’s t test. Scale Bars: 500 µm and 100 µm (amplification).

**Figure 2.**
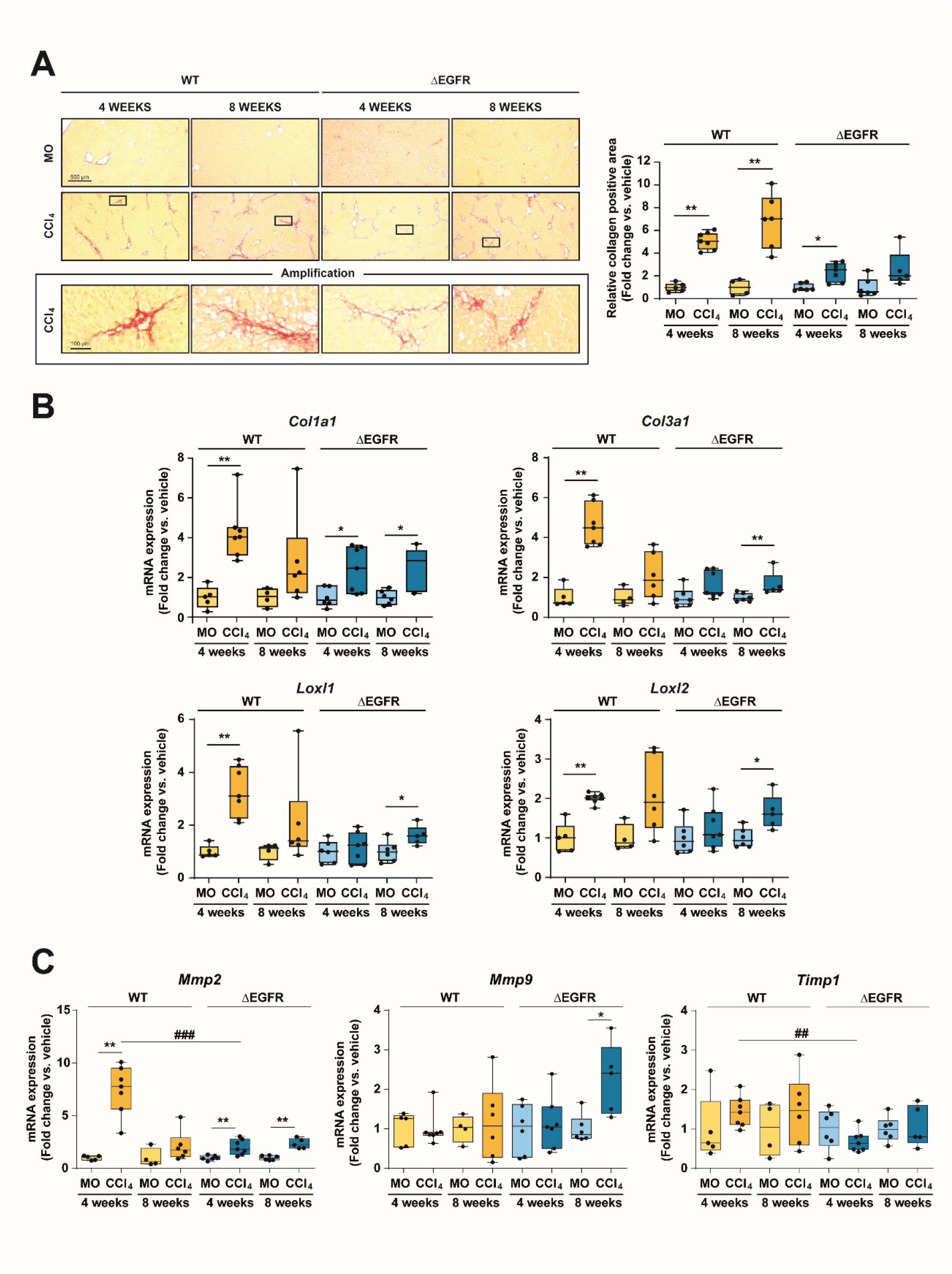
The impairment of EGFR catalytic activity in hepatocytes attenuates CCl_4_-induced liver fibrosis. **A.** Collagen deposition was assessed by Sirius red staining and positive stained area was quantified as shown on the right. **B**. RT-qPCR analysis of the hepatic mRNA expression of genes involved in extracellular matrix remodeling: *Col1a1, Col3a1, Loxl1, Loxl2, Mmp2* and *Mmp9*. Data (n=4-7 animals per group) are expressed as fold change versus vehicle (MO) and presented as box-and-whisker plots. *^,#^p<0.05, **^,##^p<0.01, ***^,###^p<0.001, using Student’s t test. * CCl4 versus MO; ^#^ ΔEGFR versus WT. Scale Bars: 500 µm and 100 µm (amplification).

### ΔEGFR livers show significant differences in the immune cell population in the CCl_4_ model of liver injury

With the aim of comparing the inflammatory response in WT and ΔEGFR mice treated with CCl_4_, we isolated and analyzed by flow cytometry the intrahepatic immune cell populations after 4 weeks of treatment. No increase in the percentage of resident macrophages was observed (***Fig. 3A***), which was corroborated by analyzing the expression of *Adgre1* and immunohistochemistry of F4/80 in tissue sections (***Suppl. Fig. 3***). However, ΔEGFR livers presented a decrease in the late recruited macrophages (correlating with the less parenchymal damage observed in Fig. 1) and the most significant result was the shift in the M1/M2 balance towards M2 polarity (***Fig. 3C***), which correlated with a significant lower ratio between the expression of the pro-inflammatory cytokine *Il12*, characteristic of M1 macrophages, and the anti-inflammatory cytokine *Il10*, characteristic of M2 polarization, (***Fig. 3D***) detected in ΔEGFR livers. Regarding lymphocyte populations, a shift in the Cd4/Cd8 ratio was observed in favor of Cd4 lymphocytes in ΔEGFR livers (***Fig. 4A***). Moreover, this response in ΔEGFR livers concurred with significant lower levels of naïve and IL17+ Cd4+ lymphocytes, and a relevant increase in the Treg cell subpopulation (***Fig. 4B-E***).

**Figure 3.**
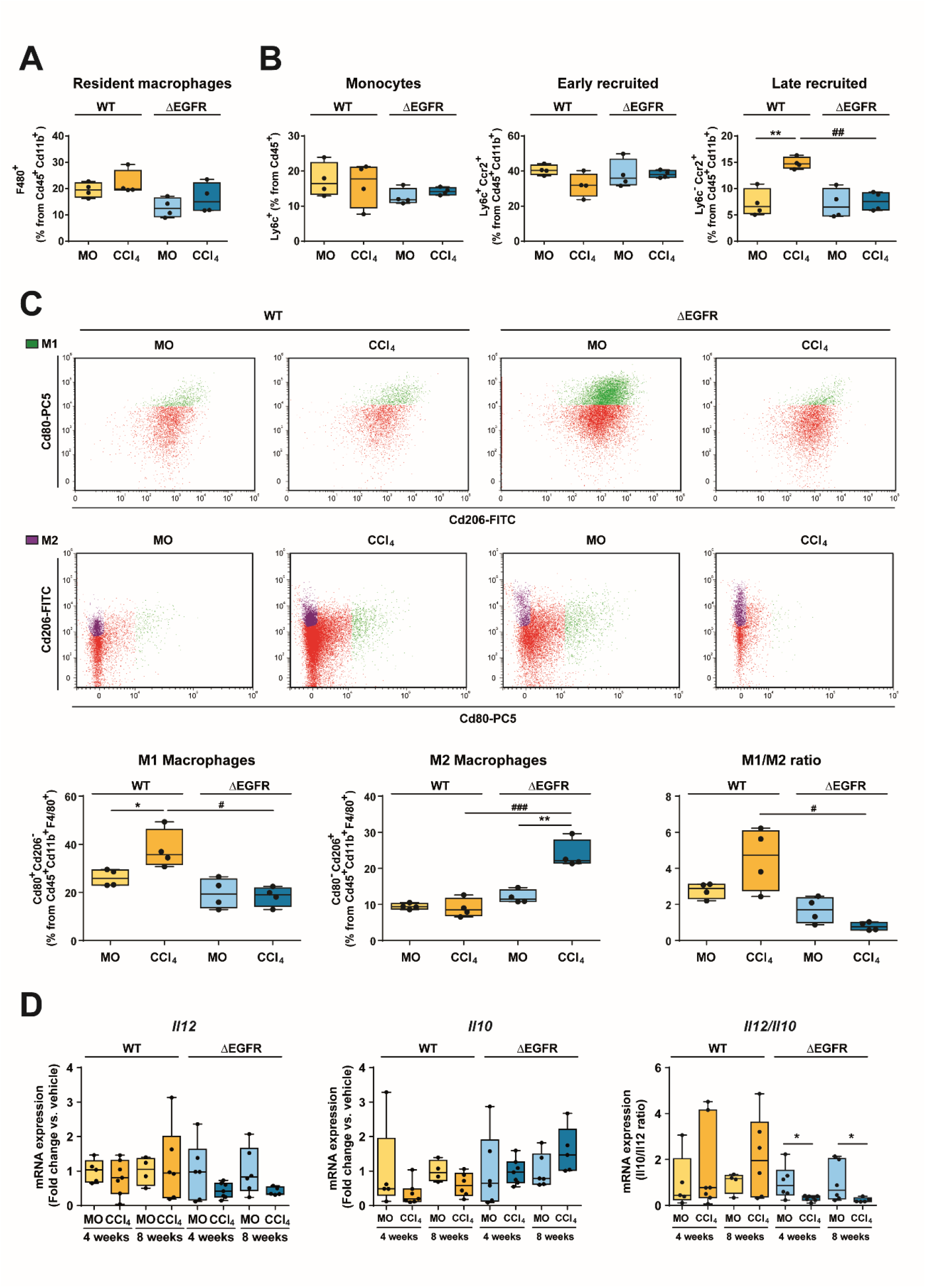
The impairment of EGFR catalytic activity in hepatocytes alters macrophages phenotype following CCl_4_-induced liver damage. **A-C:** Non-parenchymal liver cells were isolated and analyzed by flow cytometry in WT and ΔEGFR mice after 4 weeks of treatment with CCl_4_ (n=4 animals per group). **A.** Resident macrophages (percentage of F4/80+ pre-gated on Cd45+ and Cd11b+ cells). **B.** Monocytes (percentage of Ly6c+ pre-gated on Cd45+ cells) and early macrophages (percentage of Ly6c+ Ccr2+ pre-gated on Cd45+ and Cd11b+ cells) and late recruited macrophages (percentage of Ly6c-Ccr2+ pre-gated on Cd45+ and Cd11b+ cells). **C.** M1 (percentage of Cd80+ Cd206+ pre-gated on Cd45+, Cd11b+ and F4/80+ cells) and M2 (percentage of Cd80-Cd206+ pre-gated on Cd45+, Cd11b+ and F4/80+ cells) macrophage populations (M1/M2 ratio on the right). **D.** RT-qPCR analysis of the hepatic mRNA expression of *Il10* and *Il12* mRNA levels (fold change). *Il12/Il10* ratio (on the right) was used as indicator of the pro/anti-inflammatory phenotype of macrophages (n=4-7 animals per group). Data are presented as box-and-whisker plots. *^,#^p<0.05, **^,##^p<0.01, ***^,###^p<0.001, using Student’s t test. * CCl_4_ versus vehicle (MO); ^#^ ΔEGFR versus WT.

**Figure 4.**
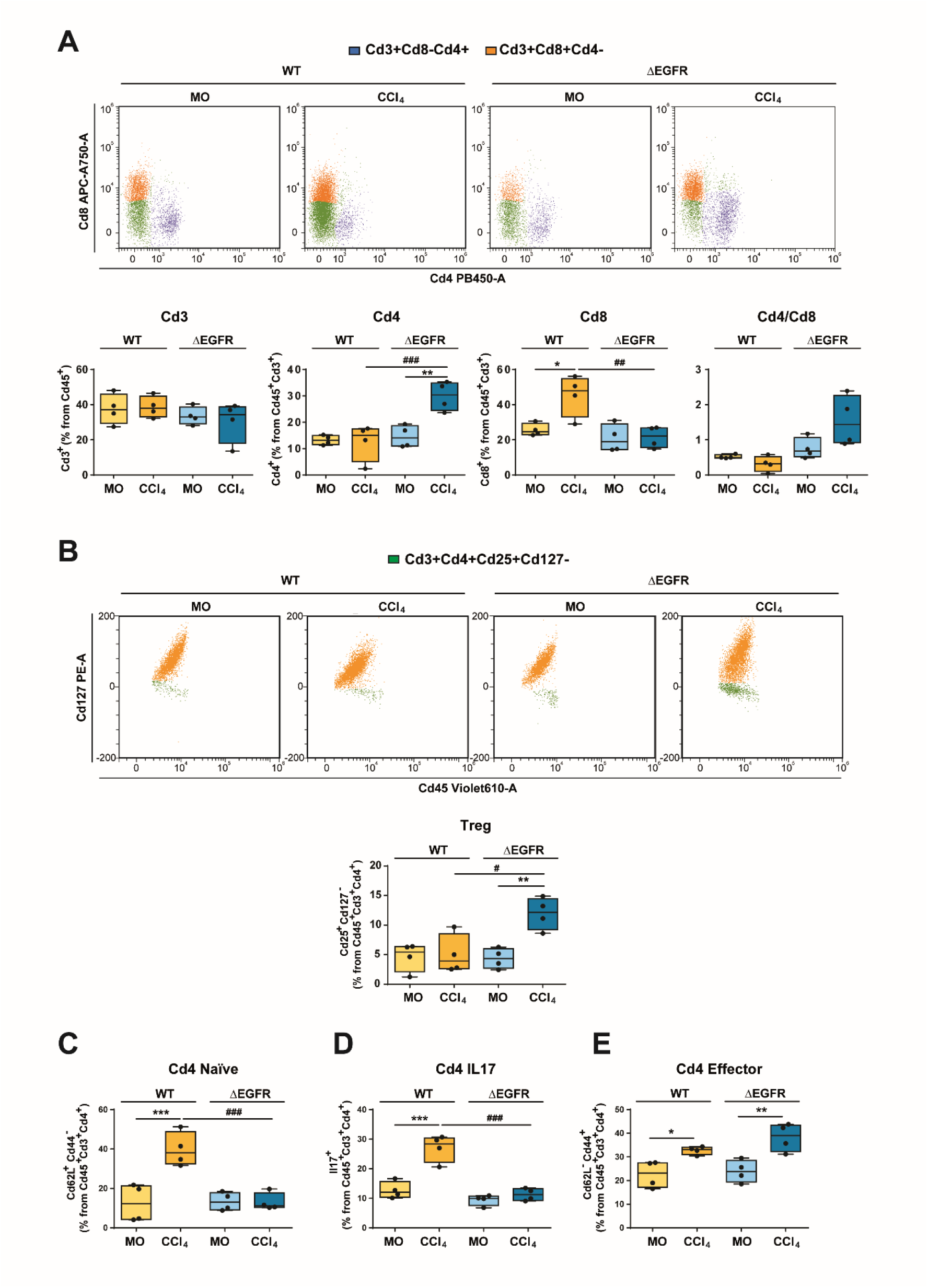
The impairment of EGFR catalytic activity in hepatocytes alters the lymphocytic infiltrate following CCl_4_-induced liver damage. Non-parenchymal liver cells were isolated and analyzed by flow cytometry in WT and ΔEGFR mice after 4 weeks of treatment with CCl_4_ (n=4 animals per group). **A.** Cd4+ (percentage of Cd4+ pre-gated on Cd45+ and Cd3+ cells) and Cd8+ (percentage Cd8+ pre-gated on Cd45+ and Cd3+ cells) cell populations (Cd4/Cd8 ratio on the right). **B.** Changes in Cd4 lymphocytes subpopulations: Treg (percentage of Cd25+ Cd127-pre-gated on Cd45+ Cd3+ and Cd4+ cells) ; **C.** Cd4 Naïve (percentage of Cd62L+ Cd44-pre-gated on Cd45+ Cd3+ and Cd4+ cells); **D.** Cd4 IL17 (percentage of Il17+ pre-gated on Cd45+ Cd3+ and Cd4+ cells) ; **E.** Cd4 Effector (percentage of Cd62L-Cd44+ pre-gated on Cd45+ Cd3+ and Cd4+ cells). Data are presented as box-and-whisker plots. *^,#^p<0.05, **^,##^p<0.01, ***^,###^p<0.001 using Student’s t test. * CCl_4_ versus vehicle (MO); ^#^ ΔEGFR versus WT.

### The EGFR catalytic activity regulates the hepatocyte gene transcriptome in response to CCl_4_-induced liver damage

To deepen into the underlying molecular mechanisms by which the EGFR pathway in hepatocytes could be controlling the fibrotic niche interactome, we next performed RNA-seq analysis in hepatocytes isolated from WT and ΔEGFR mice either treated with mineral oil or CCl_4_ for 4 weeks (***Suppl. Fig. 4A***). The most relevant results are summarized in the ***Fig. 5***. When compared the changes in gene expression caused by CCl_4_ treatment versus control samples in both WT and ΔEGFR hepatocytes, 1258 genes were specifically differentially expressed in the WT comparison, whereas 209 genes were only differentially expressed in the one from EGR. (***Fig. 5A***). Transcription factor activity was inspected by functional enrichment analysis, based on the differentially expressed genes of each comparison. Treated hepatocytes from WT mice showed 265 TF binding sites significantly enriched among those genes, which did not appear in EGFR mice (***Fig. 5B*** and details in ***Suppl. Fig. 4B***). Many of the TF whose activity appeared modulated in WT hepatocytes were related to proliferation and differentiation (EGR, Myc, E2F, Ets-related, KLFs, etc.), stress pathways (HIF-1, p53) and metabolism (PPARa, SREBP, CREB-related). The most significant difference in the ΔEGFR hepatocytes was the increase in the activity of PPARg, as well as GLI or NF1. In addition to the expected differences in the activation of the EGFR pathway (***Suppl. Fig. 5A***), relevant differences on tissue remodeling, collagen metabolic processes or regulation of inflammation and cytokine production pathways were identified in hepatocytes from WT but not from ΔEGFR mice (***Fig. 5C***). The cellular response to xenobiotic stimulus was activated in WT and ΔEGFR mice (***Suppl. Fig. 5A***), which would indicate that the response to CCl_4_ has not been affected by silencing EGFR in hepatocytes.

**Figure 5.**
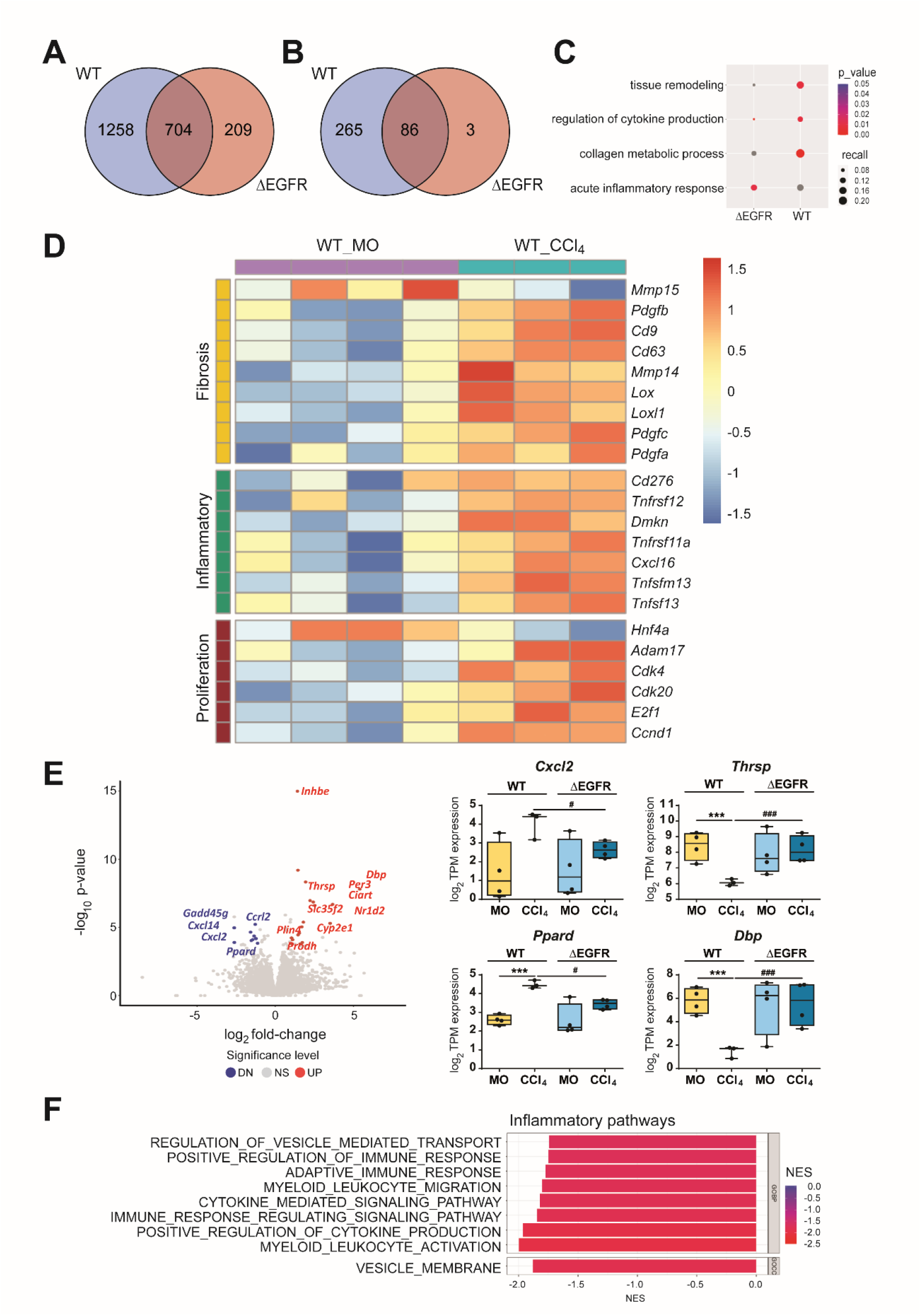
The EGFR pathway regulates the hepatocyte gene transcriptome during the response to CCl_4_. RNA-seq analysis in hepatocytes isolated from WT and ΔEGFR mice treated with either vehicle (mineral oil, MO) or CCl_4_ for 4 weeks (4 animals per group, excepting WT-CCl_4_, which were 3). **A.** Venn diagram, comparison of differentially expressed genes (DEG) detected in CCl4 vs vehicle for WT and ΔEGFR mice. **B.** Venn diagram, comparison of transcription factor binding sites found enriched by gProfiler for both contrasts. **C.** Dot plot showing differences in enrichment for key events in chronic liver damage between ΔEGFR and WT mice. **D.** Heatmap showing changes in the expression of genes related to fibrosis, inflammation or proliferation that appeared specifically in WT, but not in ΔEGFR, hepatocytes after CCl_4_ treatment. **E. Left:** Volcano plot showing differences in the expression of genes in ΔEGFR versus WT hepatocytes isolated after 4 weeks of CCl_4_ treatment. Differentially expressed genes (p < 0.05 and |log_2_ fold-change|> 1) are shown in red (upregulated, UP) and blue (downregulated, DN), the non-significant genes are shown in grey. **Right:** mRNA expression of *Cxcl2, Thrsp, Ppard and Dbp,* as examples of some of the genes that appeared in the Volcano plot, in hepatocytes from WT *and* ΔEGFR mice (treated with either MO or CCl_4_), analyzed by RNA-seq. **F.** Barplot of inflammatory pathways that appeared enriched when comparing treated hepatocytes, ΔEGFR versus WT, by GSEA. *^,#^p<0.05, **^,##^p<0.01, ***^,###^p<0.001. * CCl_4_ versus vehicle (MO); ^#^ ΔEGFR versus WT.

Among the genes specifically regulated in hepatocytes from CCl_4_-treated WT mice that did not appear modulated in ΔEGFR hepatocytes, we found genes related to fibrogenesis, such as members of the *Pdgf* family, metalloproteases *Mmp15* or *Mmp14* or *Lox* and *Loxl1*, among others (***Fig. 5D***), as well as genes involved in regulation of inflammation, such as cytokines/chemokines or genes related to the TNF receptor signaling (***Fig. 5D***). A Volcano plot showing differences in the expression of genes when compared ΔEGFR versus WT hepatocytes isolated after 4 weeks of CCl_4_ treatment revealed significant decrease in the expression of inflammatory genes, such as the cytokines *Cxcl2* or *Cxcl14* or the chemokine receptor *Ccrl2* (***Fig. 5E**, Left***). In fact, GO enrichment analysis also revealed that ΔEGFR cells presented a significant decrease in inflammatory pathways (***Fig. 5F***). The Volcano representation also revealed differences in genes related to differentiation, such as *Cyp2e1,* and metabolism, such as *Ppard*, *Dbp* or *Thrsp* (***Fig. 5E**, Left***). In most of the cases, CCl4 treatment induced changes in gene expression in WT hepatocytes, which were not observed in ΔEGFR hepatocytes (***Fig. 5E**, Right***). Worthy to mention as well that hepatocytes from CCl_4_-treated WT mice, but not those from ΔEGFR mice, showed relevant changes in the expression of genes related to cell cycle progression and cell proliferation (***Fig. 5D***), which would indicate higher hepatocyte regeneration in the WT than in the ΔEGFR mice. In parallel, hepatocyte identity genes, such as *Hnf4a* (***Fig. 5D***), or genes related to liver-specific functions (***Suppl. Fig. 5B***), were down-regulated. Interestingly, the ΔEGFR hepatocytes from CCl_4_-treated mice showed higher expression of mitochondrial metabolism-related pathways, such as fatty acid catabolic processes, mitochondrial respiratory chain, aerobic electron transport chain or ATP synthesis than those isolated from the WT mice (***Suppl.*** ***Fig.5C***), which indicate that they are metabolically more efficient. These metabolic changes correlated with differences in the expression and/or activity of members of the PPAR family of TF: *PPARa* or *PPARd* in WT, but *PPARg* in ΔEGFR hepatocytes (***Fig. 5*** ***and Suppl Fig. 4***). Overall, results indicate that in response to a chronic insult the EGFR pathway in hepatocytes modulates their transcriptional program.

### The EGFR catalytic activity in hepatocytes regulates the inflammatory and pro-fibrotic secretome in response to CCl_4_-induced liver damage

Analysis of the liver immune populations together with hepatocyte transcriptomic profiles seem to indicate an active participation of the hepatocyte EGFR pathway in regulating inflammation. Indeed, we next decided to explore whether secretion of factors from the hepatocytes could modulate the phenotype of macrophages. We have previously generated immortalized hepatocytes from WT and ΔEGFR mice,(10) which are a useful tool for *in vitro* approaches. Here we have analyzed how the secretome of these cells may affect the fibrotic cell interactome in *in vitro* experiments. For this, we incubated the WT and ΔEGFR hepatocytes with HB-EGF, a ligand of the EGFR, to activate the pathway (***Fig. 6A-B*)**. We next followed the experimental set up depicted in ***Fig. 6C*** to generate conditioned medium that was used to analyze the effects on the phenotype of mouse macrophages (RAW264.7 cells). As observed in ***Fig. 6D***, the conditioned medium of WT and ΔEGFR hepatocytes produced opposite effects on the expression profile of genes associated with an M1 phenotype (pro-inflammatory/fibrotic), such as *Il12*, or M2 (resolution of fibrosis), such as *Il10*. The ratio *Il12*/*Il10* significantly increased in RAW264.7 cells incubated with the conditioned medium from HB-EGF treated WT hepatocytes, but this increase was not observed when incubated with the conditioned medium from HB-EGF treated ΔEGFR hepatocytes.

**Figure 6.**
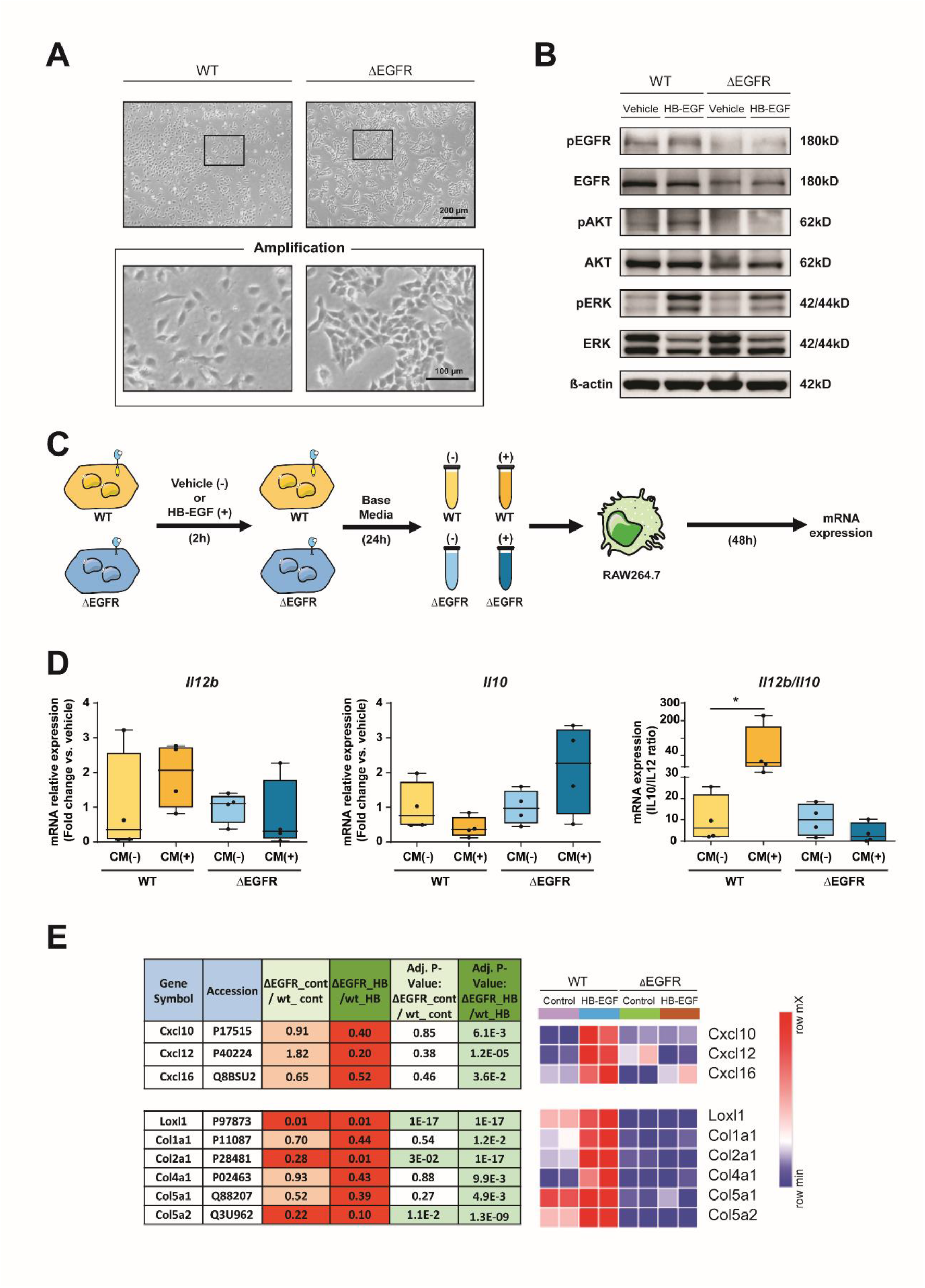
The EGFR pathway regulates the hepatocyte secretome. Effects of conditioned medium (CM) of hepatocytes from WT and ΔEGFR mice on macrophage phenotype. **A.** Representative microscopy images of cultured WT and ΔEGFR immortalized hepatocytes. **B.** Activation of key components of EGFR pathway in hepatocytes treated with the EGFR ligand HB-EGF (20 ng/mL) for 30 minutes, analysed by Western blot. **C.** Schematic representation of the experimental approach used to analyze how the hepatocyte secretome affects the phenotype of mouse macrophages (RAW264.7 cell line). **D.** Expression of M1 and M2 phenotype associated genes in macrophages, measured by RT-qPCR. *Il12/Il10* ratio was used as indicator of the pro/anti-inflammatory phenotype of macrophages. Data are expressed as fold change (CM from untreated hepatocytes, CM-, versus CM from HB-EGF-treated hepatocytes, CM+) and presented as box-and-whisker plots. **E.** Proteomic analysis of the culture medium of hepatocytes from the experiments presented in panel C. Left panel shows changes in the expression of pro-inflammatory and pro-fibrotic mediators in WT and ΔEGFR immortalized hepatocytes upon treatment with HB-EGF. Right panel shows heatmaps illustrating these results. Data from four independent experiments were analyzed using Student’s t test. *p<0.05. Scale Bars: 200 µm

To analyze whether the EGFR pathway would be also regulating the human macrophage phenotype, we used a human liver tumor cell line, the Hep3B cells, and a human monocyte cell line, the THP-1 cells. Using a similar experimental approach (***Suppl. Fig. 6A***), conditioned medium of Hep3B cells untreated or treated with HB-EGF was collected to analyze the effects on THP1 cells. Results indicated that the conditioned medium from HB-EGF treated Hep3B cells increased monocyte adhesion (***Suppl. Fig. 6B***) and the expression of pro-inflammatory genes (M1 phenotype), downregulating the expression of anti-inflammatory related genes (M2 phenotype) (***Suppl. Fig. 6C***). Interesting to observe that, again, the *IL12/Il10* ratio significantly increased in THP1 cells after the incubation with the conditioned medium from HB-EGF treated Hep3B cells ***Suppl. Fig. 6D***).

Overall, results would indicate that hepatocytes may actively participate in secreting factors that regulate the inflammatory microenvironment in the fibrotic niche. To completely demonstrate this, we next decided to perform a proteomic analysis in the conditioned medium from hepatocytes obtained following the same experimental approach presented in the Fig. 6C. The most significant results are presented in the ***Fig. 6E*** and reveal new roles for the EGFR pathway in hepatocytes on protein secretion, complementing the data observed in the transcriptomic analysis. We first focused on proteins present in the secretome that showed significant differences between ΔEGFR and WT hepatocytes. WT hepatocytes responded to HB-EGF inducing the secretion of Cxcl10, Cxcl12 and Cxcl16 and this increase was not observed in ΔEGFR hepatocytes, reinforcing the role of hepatocytes in the secretion of inflammatory cytokines under the control of the EGFR pathway. Moreover, it was also very interesting to observe that WT hepatocytes responded to HB-EGF inducing the secretion of different proteins involved in fibrosis, among them, different members of collagens, or proteins involved in their maturation, such as Loxl1. ΔEGFR hepatocytes secreted much lower levels of all these proteins and as expected, no change was observed in response to HB-EGF. Indeed, the secretome analysis also revealed a role for hepatocytes in the production of ECM proteins.

### The EGFR pathway is activated during the progression of fibrosis in humans, correlating with expression of specific fibrotic and inflammatory genes

To analyze the translational relevance of the results presented above, data from gene expression in a cohort of liver biopsies from HCC-naïve NAFLD patients was accessed through GEO accession number GSE193066 (see Material and Methods section). We could observe that the relative enrichment of EGF/EGFR signaling (WikiPathways) was significantly increased during progression of liver fibrosis (***Fig. 7A***). The relative enrichment of EGFR target genes (Gene Transcription Regulation Database) also presented a tendency, almost significant, to increase at advanced stages of fibrosis. Interestingly, the expression of fibrosis-related genes, such as collagen genes (*COL1A1*, *COL3A1*, *COL4A1*, *COL5A1*), *LOXL1*, *MMP2*, *MMP9* or *TIMP1* presented a significant positive correlation with the EGF/EGFR signaling (***Fig. 7B***). Regarding genes related to the immune system, it was relevant that *CXCL10*, which was found regulated by the EGFR pathway in the secretome analysis in hepatocytes (***Fig. 6E***), showed increased expression at advanced stages of fibrosis (***Fig. 7C***), and the expression of *CXCL10* and *CXCL12* correlated with the EGF/EGFR signaling (***Fig. 7D***). Finally, a differential pattern was found in the expression of *CD4* and *CD8A*. The expression of *CD4* decreased, whereas the expression of *CD8* increased at advanced stages of fibrosis (***Fig. 7D***), which correlated with the differential pattern of Cd8+ and Cd4+ lymphocytes found in the ΔEGFR livers after CCl_4_ treatment (***Fig. 4***). Furthermore, the expression of CD8A positively correlated with the EGF/EGFR signaling (***Fig. 7E***). These data indicate that the results found in the mice model may help to better understand the fibrosis process in humans.

**Figure 7.**
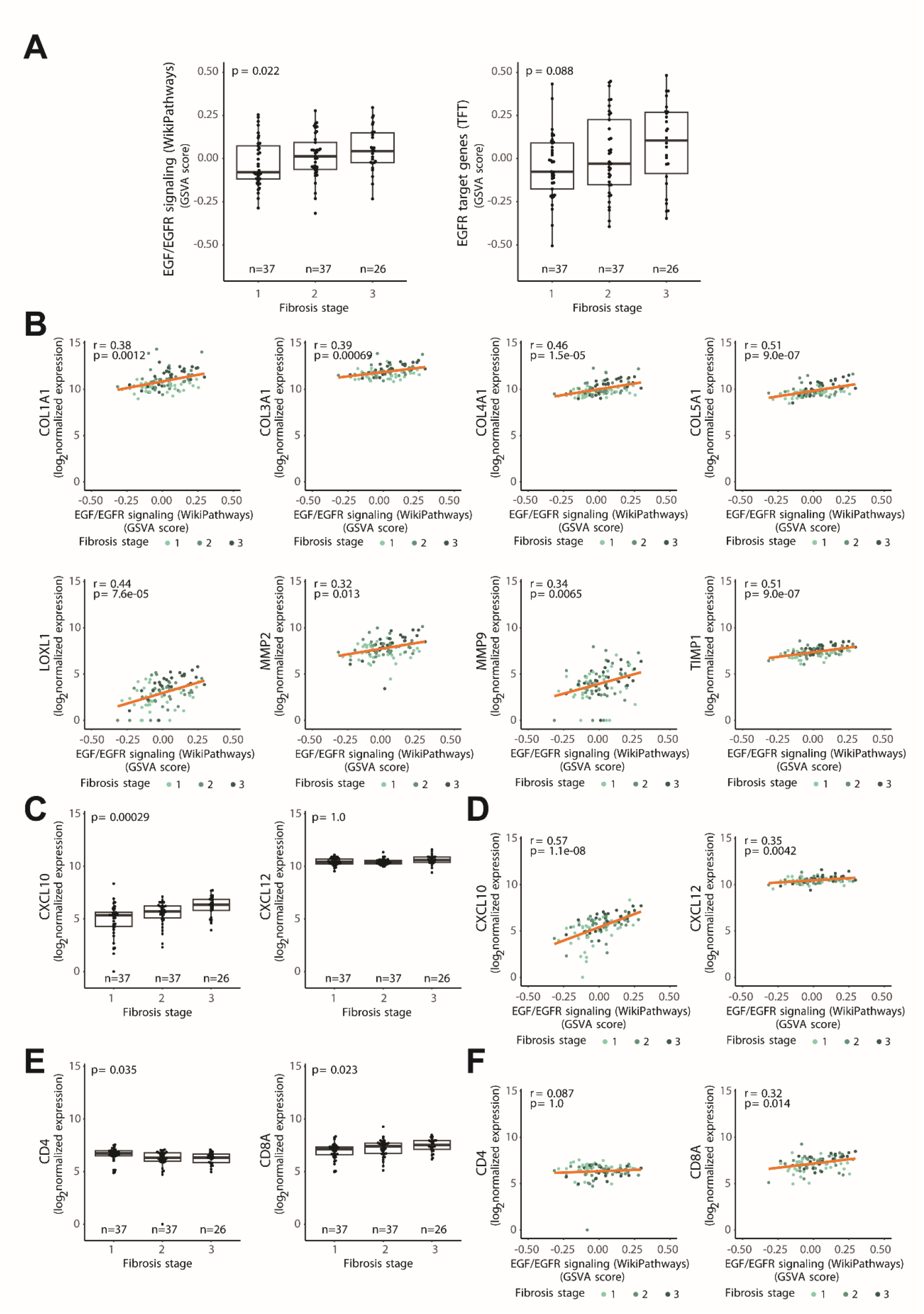
The EGFR pathway is activated in human fibrosis. **A.** Boxplots of relative enrichment (GSVA score) of either EGF/EGFR signaling or EGFR target genes across fibrosis stages. Positive values indicate a higher relative activation of the gene set while lower values indicate a relative lower activation of the pathway. **B.** Pearson correlation of fibrosis-related genes expression with the relative enrichment of EGF/EGFR signaling. Each dot is a sample, and its color indicate the fibrosis stage. **C**. Boxplot of *CXCL10* and *CXCL12* across fibrosis stages. **D**. Pearson correlation between the relative enrichment of EGF/EGFR signaling and *CXCL10* and *CXCL12* gene expression. Each dot is a sample, and its color indicate the fibrosis stage. **E**. Boxplot of *Cd4* and *Cd8A* across fibrosis stages. **F**. Pearson correlation between the relative enrichment of EGF/EGFR signaling and *Cd4* and *Cd8A* gene expression. Each dot is a sample, and its color indicate the fibrosis stage. Data was analyzed using Kendall’s tau test and all p-values were adjusted for multiple testing.

## DISCUSSION

Development of liver fibrosis is frequently the consequence of hepatocyte chronic damage, HSC activation and proliferation, and changes in ECM production and accumulation.(1) This process is accompanied by amplified inflammatory responses, with immune cells, especially macrophages, recruited to the site of injury and activated in order to orchestrate the process of wound healing and tissue repair.(25, 26) Unidentified fibrosis can evolve into more severe consequences over the time, such as cirrhosis and liver cancer.(27) Improving the knowledge about the intercellular interactions in the fibrotic niche is necessary to better understand this process and uncover potential therapeutic targets. In response to injury, hepatocytes change their gene expression and secretion profile. Although some of these newly expressed factors have been identified as fibrogenic,(1) how the hepatocyte gene and secretome signatures may regulate the advance in chronic liver injury and inflammation is poorly understood yet. This work demonstrates that the EGFR pathway in hepatocytes is required for an efficient progression of the fibrotic process in response to CCl_4_ treatment in mice. Attenuation of the EGFR kinase activity in hepatocytes reduces the parenchymal liver damage and the extension of fibrotic areas, correlating with decrease in the expression of collagens and Loxl genes, particularly *Loxl1*, increase in *Mmp9* and decrease in *Timp1*. Previous studies had supported a role for the EGFR1 signaling pathway in the initiation and development of liver fibrosis(28) and pharmacological inhibition of EGFR attenuates liver fibrosis in some mice experimental models of liver fibrosis, including non-alcoholic fatty liver disease (NASH).(5–7, 29, 30) However, in most of these works, the attention was mainly focused on the role of the EGFR pathway on hepatic steatosis and/or on HSC/myofibroblast activation and proliferation. The main work that analyzed the role of the hepatocyte ERBB tyrosine kinases comes from Dr. Russell’s group, which suggested that EGFR-ERBB3 heterodimeric signaling in damaged hepatocytes may play a relevant role in liver fibrosis.(9) Our study constitutes a step forward for deepening into the molecular mechanisms that are regulated by the EGFR in hepatocytes during chronic liver damage.

We first focused on inflammation since previous studies from our group in these ΔEGFR mice had demonstrated that EGFR catalytic activity is critical for generating an efficient inflammatory ΔEGFR mice process during DEN-induced tumorigenesis.(12) In a more recent study, we also found that after chronic feeding with 3,5-diethoxycarbonyl-1,4-dihydrocollidine (DDC), which induced cholestatic damage, ΔEGFR mice presented a reduced and delayed liver damage and a more efficient regeneration upon DDC injury, concomitantly with a shift from a profibrotic to a restorative inflammatory response, an enhanced ductular reaction and expansion of HPC/oval cells.(31) Results here shown strongly indicate that after CCl_4_ treatment, ΔEGFR livers show significant differences in the immune landscape, favoring the anti-inflammatory (M2) phenotype in macrophages. Despite tremendous progress, numerous challenges remain in deciphering the full spectrum of macrophage activation and its implication in either promoting liver disease progression or repairing injured liver tissue.(32) While in homeostatic conditions resident macrophages can be identified in the liver, in recent years numerous studies have provided compelling evidence that in inflammatory conditions there is considerably more heterogeneity among hepatic macrophages.(33, 34) Experimental animal models indicate that macrophages are critical for fibrosis regression because they play key roles in repairing the necrotic lesions, not only by removing necrotic tissues, but also by inducing cell death-resistant hepatocytes, activating α-smooth muscle actin-expressing HSCs to facilitate necrotic lesion resolution, and exerting anti-inflammatory actions.(35, 36) Although the role of M2-like macrophages in liver fibrosis is controversial yet, strong evidence supports that they could exert protection against the injury, among other actions, by inhibiting necroptosis and conferring apoptosis resistance to hepatocytes.(37, 38) Furthermore, M2 macrophage-derived exosomes significantly inhibit HSCs activation.(39) In addition to the observed effects on macrophage phenotype, the attenuation of the EGFR catalytic activity in hepatocytes provoked notable differences in the T cell populations, with an increase in the Treg lymphocytes, as well as significant differences in the proportion of Cd8+ and Cd4+ cells, decreasing the Naïve and IL17+ cells. Th17/Treg imbalance exists in mice with liver fibrosis, which potentially promotes liver fibrosis via HSC activation.(40) The balance between Th17 and Treg cells is critical for immune reactions, especially in injured liver tissue and the return to immune homeostasis.(41) It has been proposed that by inhibiting profibrotic Th17 and Cd8+ T cells, Treg cells may help to maintain a healthy inflammatory environment, regulating the aberrant activation and functions of immune effector cells.(42) Overall, results strongly indicate that the activation of the EGFR pathway in hepatocytes during chronic liver injury, may have consequences on the inflammatory environment, regulating macrophage plasticity and favoring a high ratio Th17/Treg cells.

Analysis of the transcriptional signature of hepatocytes from fibrotic WT and ΔEGFR livers revealed the role of the EGFR pathway in regulating the expression of genes that promote fibrogenesis, such as different forms of the *Pdgf* gene, as well as genes coding for lysyl oxidases, such as *Lox* and *Loxl1*, involved in collagen chain trimerization. Different genes related to inflammatory processes, such as genes related to the TNF pathway (*Tnfrsf12, Tnfrf11a, Tnfsfm13*), the proinflammatory dermokine (*Dmkn*), or regulators of the T cell mediated response, such as *Cd276*, appeared also upregulated specifically in WT livers from CCl_4_-treated mice. Overall, the EGFR pathway modulates a transcriptomic program in hepatocytes that contribute to generate a pro-inflammatory and fibrotic environment, which justifies the strong differences observed between WT and ΔEGFR mice after CCl_4_ treatment. Correlating with this, numerous TF appeared regulated in CCl_4_-treated WT but not in ΔEGFR hepatocytes. An aspect that caught our attention was the strong differences in the expression of genes related to proliferation and differentiation, which were up-regulated and down-regulated, respectively, in WT but not in ΔEGFR hepatocytes after CCl_4_ treatment. Transient hepatocellular dedifferentiation can occur as part of the regenerative mechanisms triggered in response to acute liver injury. However, persistent downregulation of key identity genes is now accepted as a strong determinant of organ dysfunction in chronic liver disease,(43) by stably reprogramming the critical balance of transcription factor responsible for hepatocyte identity, among them, HNF4α.(43, 44) Hepatocytes from CCl_4_-treated ΔEGFR hepatocytes maintain *Hnf4a* expression and do not show changes in expression of proliferation-differentiation genes, which may suggest that damage of hepatocytes is much lower and/or that the EGFR pathway is required for this gene transcription reprogramming. In parallel, ΔEGFR hepatocytes display higher expression of mitochondrial metabolism pathways-related genes which might confer them higher efficient energy capacity.

Since transcriptomic analysis revealed differences in genes coding for cytokines or chemokines, as well as exosomal markers, such as *Cd9* or *Cd63*,(45) we decided to analyze the differences in the secretome of hepatocytes of WT and ΔEGFR hepatocytes either untreated or treated with an EGFR ligand, HB-EGF. Results demonstrated that the activation of the EGFR pathway conferred hepatocytes’ secretome the capacity to modulate macrophages polarity promoting an anti-inflammatory phenotype. The proteomic analysis of the conditioned media revealed that the secretion of different chemokines, such as Cxcl10, Cxcl12 or Cxcl16, were significantly increased by HB-EGF in WT hepatocytes, but not in ΔEGFR hepatocytes. These cytokines have been previously related to the progression of chronic liver injury. Intrahepatic CXCL10 has been strongly associated to liver fibrosis in different context in human liver pathologies(46, 47) through regulation of macrophage-associated inflammation(48) or the infiltration of immune T cells.(46) CXCL12, the ligand of CXCR4, can be involved in multiple pathological mechanisms in fibrosis, such as inflammation, immunity, epithelial-mesenchymal transition, and angiogenesis.(49) Furthermore, the secretome analysis reflected a role for hepatocytes in producing ECM proteins under the control of the EGFR pathway. Worthy to highlight that Loxl1 appeared in all the studies presented in this manuscript: in the *in vivo* experiments, in the transcriptomic analysis performed in hepatocytes, as well in the secretome. It is very clear that hepatocytes contribute to the expression, production, and secretion of lysyl oxidases required for the final formation of the collagen fibers, and the EGFR pathway regulates this process.

Our data reveal that the findings obtained in the mouse model could be translated to human patients suffering a fibrotic process. The analysis of transcriptomic data in a cohort of liver biopsies from HCC-naïve NAFLD patients evidenced that the EGFR signaling pathway increases in advanced stages of fibrosis and correlates with the expression of fibrotic and inflammatory genes identified in the *in vivo* and *in vitro* experiments. Expression of *Loxl1*, mentioned above as a clear candidate to be under the control of the EGFR pathway, as well as *Timp1*, which has been also identified in some of the experimental analyses in the mice model, showed a strong correlation with the EGF/EGFR signaling pathway. Also, it is worth highlighting *Cxcl10* that shows a strong increase in advanced stages of fibrosis, correlating with the EGF/EGFR signaling, or *CD4 and CD8A*, presenting inversed correlations with fibrosis stage. Altogether, data in the human samples may indicate that genes that play essential roles during liver fibrosis/inflammation may be under the control of the EGFR pathway.

Concluding, data presented here led us to propose a model where the EGFR pathway in hepatocytes would control liver fibrosis and inflammation in response to a chronic insult on the liver through remodeling the hepatocyte gene transcriptome/secretome, which has consequences on the deposit of ECM as well as on the immune microenvironment. Overall, our study uncovers novel mechanistic insights on EGFR kinase-dependent actions in hepatocytes that reveals its key role in chronic liver damage.

## Supporting information

Visual Abstract

## Acknowledgments

Authors are very grateful to Drs. Jose Carlos Segovia and Lluis Montoliu for their initial contribution in the generation of the transgenic mice. We also thank the technical assistance from UCM Proteomics Unit for the proteomic analysis performed in hepatocytes conditioned medium.

