## Supplementary Material for "The hepatocyte Epidermal Growth Factor Receptor (EGFR) pathway regulates the cellular interactome within the liver fibrotic niche"

**Supplementary Table 1. Primary antibodies used for immunodetection.**

| Protein | Antibody | Company | Type | Source | Use | Dilution |
|---|---|---|---|---|---|---|
| α-SMA | ab5694 | Abcam | Polyclonal | Rabbit | WB | 1/2000 |
| | | | | | IHC | 1/50 |
| F4/80 | Ab6640 | Abcam | Monoclonal | Rat | IHC | 1/50 |
| pSMAD3 | 07-1389 | Millipore | Polyclonal | Rabbit | WB | 1/1000 |
| SMAD3 | 04-1035 | Millipore | Monoclonal | Rabbit | WB | 1/1000 |
| pEGFR | CST 3777S | CST | Monoclonal | Rabbit | WB | 1/500 |
| EGFR | CST 2232S | CST | Polyclonal | Rabbit | WB | 1/1000 |
| pAkt | CST 4060 | CST | Monoclonal | Rabbit | WB | 1/1000 |
| Akt | CST 9272 | CST | Polyclonal | Rabbit | WB | 1/1000 |
| pERK | CST 9101 | CST | Polyclonal | Rabbit | WB | 1/1000 |
| ERK | CST 4695 | CST | Monoclonal | Rabbit | WB | 1/1000 |
| β-actin | A5441 | Sigma-Aldrich | Monoclonal | Mouse | WB | 1/5000 |

WB, Western Blot; IHC, immunohistochemistry.

**Supplementary Table 2. Antibodies used for immunodetection of liver immune populations by FACS.**

| | Company | Type | Source | Dilution |
|---|---|---|---|---|
| FITC Rat Anti-Mouse Cd45. Clone I3/2.3 | Southern Biotech | IgG | Rat | 1/100 |
| APC F4/80 Monoclonal Antibody. Clone BM8 | eBioscience | IgG2ak | Rat | 1/50 |
| PE/Cyanine7 anti-mouse Cd11b(Mac1) Clone M1/70 | BioLegend | IgG2bk | Rat | 1/100 |
| PE/Cyanine7 anti-mouse Cd3. Clone 145-2C11 | BioLegend | IgG | Hamster | 1/100 |
| PE Rat Anti-Mouse Cd8. Clone 53-6.7 | Beckman Coulter | IgG2ak | Rat | 1/50 |
| PerCP/Cyanine5.5 Anti-mouse Cd4. Clone GK1.5 | BioLegend | IgG2bk | Rat | 1/50 |
| PE anti-mouse Cd206. Clone C068C2 | BioLegend | IgG2ak | Rat | 1/25 |
| FITC anti-mouse CD25. Clone 7D4 | Pharmingen | IgMk | Rat | 1/25 |
| PE anti-mouse Cd127. Clone A7R34 | eBioscience | IgG2ak | Rat | 1/25 |
| Brilliant Violet 570 Anti-mouse Cd45. Clone 30-F11 | BioLegend | IgG2b | Rat | 1/100 |
| PE anti-mouse IL-17A. Clone TC11-18H10.1 | BioLegend | IgG1k | Rat | 1/25 |
| APC anti-mouse Cd62L. Clone MEL-14 | eBioscience | IgG2ak | Rat | 1/25 |
| PECy5 anti-mouse Cd44 (SRPD). Clone IM7 | BioLegend | IgG1k | Rat | 1/25 |

**Supplementary Table 3. Primers used for RT-qPCR (mouse genes).**

| Gene | Forward | Reverse |
|---|---|---|
| *Col1a1* | GAGAGGTGAACAAGGTCCCG | AAACCTCTCTCGCCTCTTGC |
| *Col3a1* | GACCAAAAGGTGATGCTGGACAG | CAAGACCTCGTGCTCCAGTTAG |
| *Loxl1* | GAGTGCTATTGCGCTTCCC | GGTTGCCGAAGTCACAGGT |
| *Loxl2* | TTCTGCCTGGAGGACACTGAGT | TCGGTGATGTCTATCCACTGGC |
| *Mmp2* | GTGGGACAAGAACCAGATCAC | GCATCATCCACGGTTTCAG |
| *Mmp9* | CCTGGCTCTCCTGGCTTT | AGCGGTACAAGTATGCCTCTG |
| *Timp1* | TGGCATCCTCTTGTTGCTATCACTG | TGAATTTAGCCCTTATGACCAGGTCC |
| *Tgfb1* | GTCAGACATTCGGGAAGCAG | GCGTATCAGTGGGGGTCA |
| *Tgfb2* | TCCCCTCCGAAAATGCCATC | GGAAGACCCTGAACTCTGCC |
| *Tgfb3* | TTTGCGGAGGACGGAGTAAC | ACAGTCACCAGCATCTCAGC |
| *Adgre1* | TGCACTGACACCACAGACAG | TGCAGACTGAGTTAGGACCAC |
| *Il10* | CCTTCAGCCAGGTGAAGACT | GGCAACCCAAGTAACCCTTA |
| *Il12b* | ATTACTCCGGACGGTTCACG | ACGCCATTCCACATGTCACT |
| *Rpl32* | ACAATGTCAAGGAGCTGGAG | TTGGGATTGGTGACTCTGATG |

**Supplementary Table 4. Primers used for RT-qPCR (human genes).**

| Gene | Forward | Reverse |
|---|---|---|
| *ERBB1* (EGFR) | GCAAATTCCGAGACGAAGCC | CTGTATTTGCCCTCGGGGTT |
| *GPR18* | ACCCAAAGTCAAGGAGAAGTC | GCATCAGGAAAGCGAAACAG |
| *IL6* | ACCCCCAATAAATATAGGACTGGA | TTCTCTTTCGTTCCCGGTGG |
| *IL12B* | CTTGGACCAGAGCAGTGAGG | GAACCTCGCCTCCTTTGTGA |
| *MRC1* | GCAAAGTGGATTACGTGTCTTG | CTGTTATGTCGCTGGCAAATG |
| *EGR2* | TTGACCAGATGAACGGAGTG | GCCCATGTAAGTGAAGGTCTG |
| *IL10* | TGCCTTCAGCAGAGTGAAGA | GCAACCCAGGTAACCCTTAAA |
| *RPL32* | AACGTCAAGGAGCTGGAAG | GGGTTGGTGACTCTGATGG |

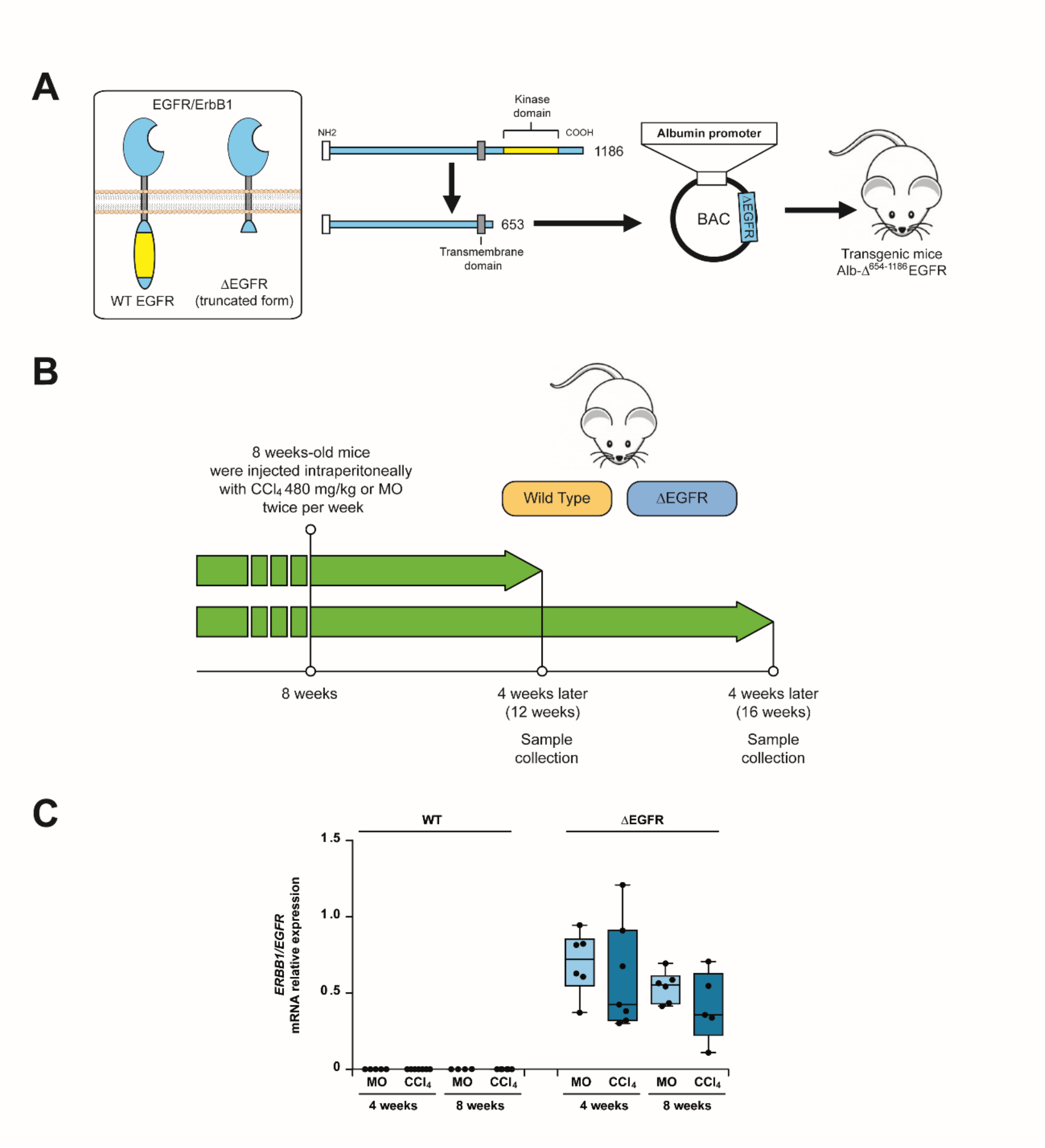


**Supplementary Figure 1. Experimental mouse model. A.** Transgenic model Alb-Δ654-1186EGFR (ΔEGFR) mice. A complementary DNA coding for a truncated form of the EGFR with a deletion in its intracytosolic region (amino acids 654-1186) was cloned in a transference plasmid under the control of the albumin promoter locus upstream ATG of the mouse albumin gene. **B.** Experimental model of $CCl_4$-induced liver fibrosis in ΔEGFR and WT mice. **C.** Hepatic mRNA expression of human *ERBB1/EGFR* was determined by RT-qPCR in livers from ΔEGFR and WT mice treated with $CCl_4$ or vehicle (mineral oil: MO) for 4 or 8 weeks. Data (n=4-7 animals per group) were analyzed using Student's t test and presented as box-and- whisker plots.

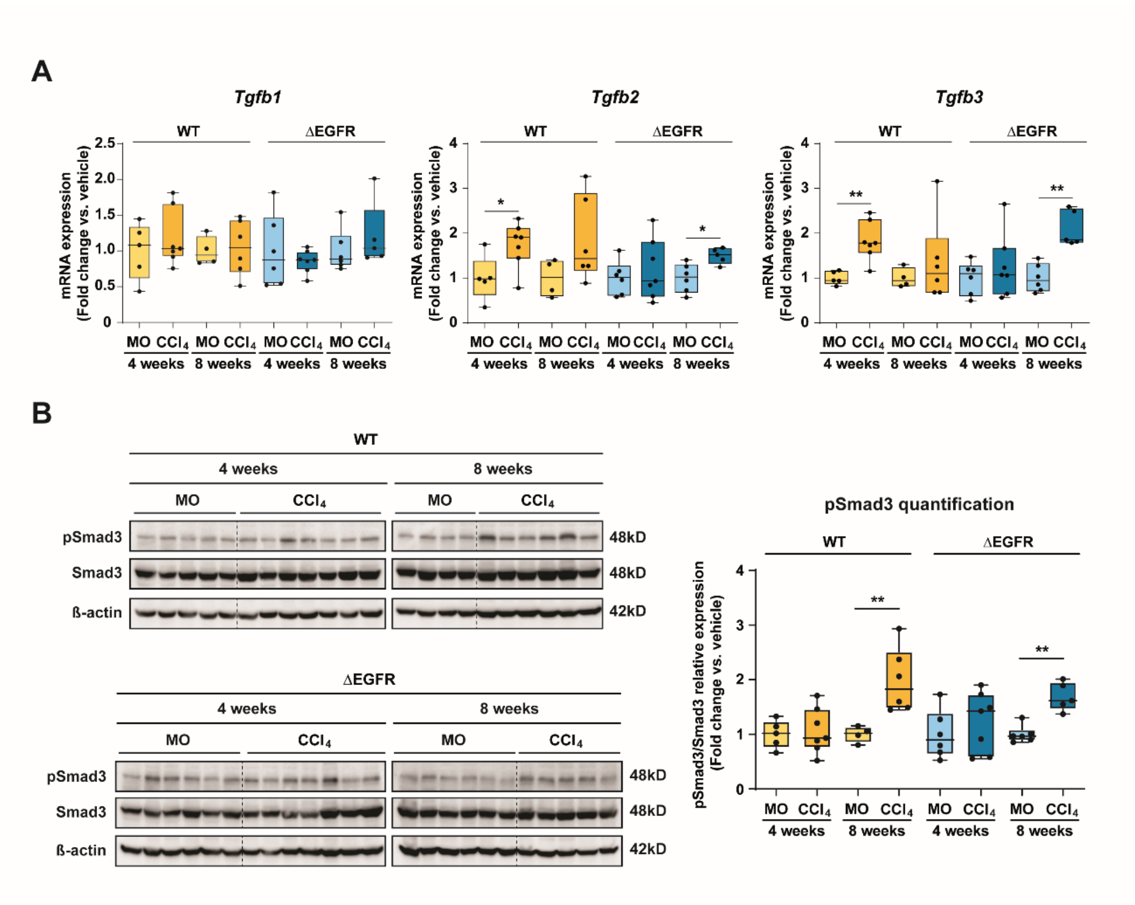


**Supplementary Figure 2. Analysis of the Transforming Growth Factor-beta (TGF-β) pathway in livers from $CCl_4$-treated ΔEGFR and WT mice.** ΔEGFR and WT mice were treated with $CCl_4$ or vehicle (MO) for 4 or 8 weeks**. A.** The mRNA expression of TGF-β ligands 1-3 (*Tgfb1, Tgfb2* and *Tgfb3*) was assessed by RT-qPCR. **B.** The activation of the signaling pathway was determined by western blot analysis of the phosphorylated Smad3 (pSmad3) protein levels (left panel). Densitometric analysis of western blot expressed as the ratio pSmad3 versus Smad3 after normalization with β-actin levels (right panel). Data (n=4-7 animals per group) are expressed as fold change versus vehicle (MO) and presented as box-and- whisker plots. *p<0.05, **p<0.01 using Student's t test.

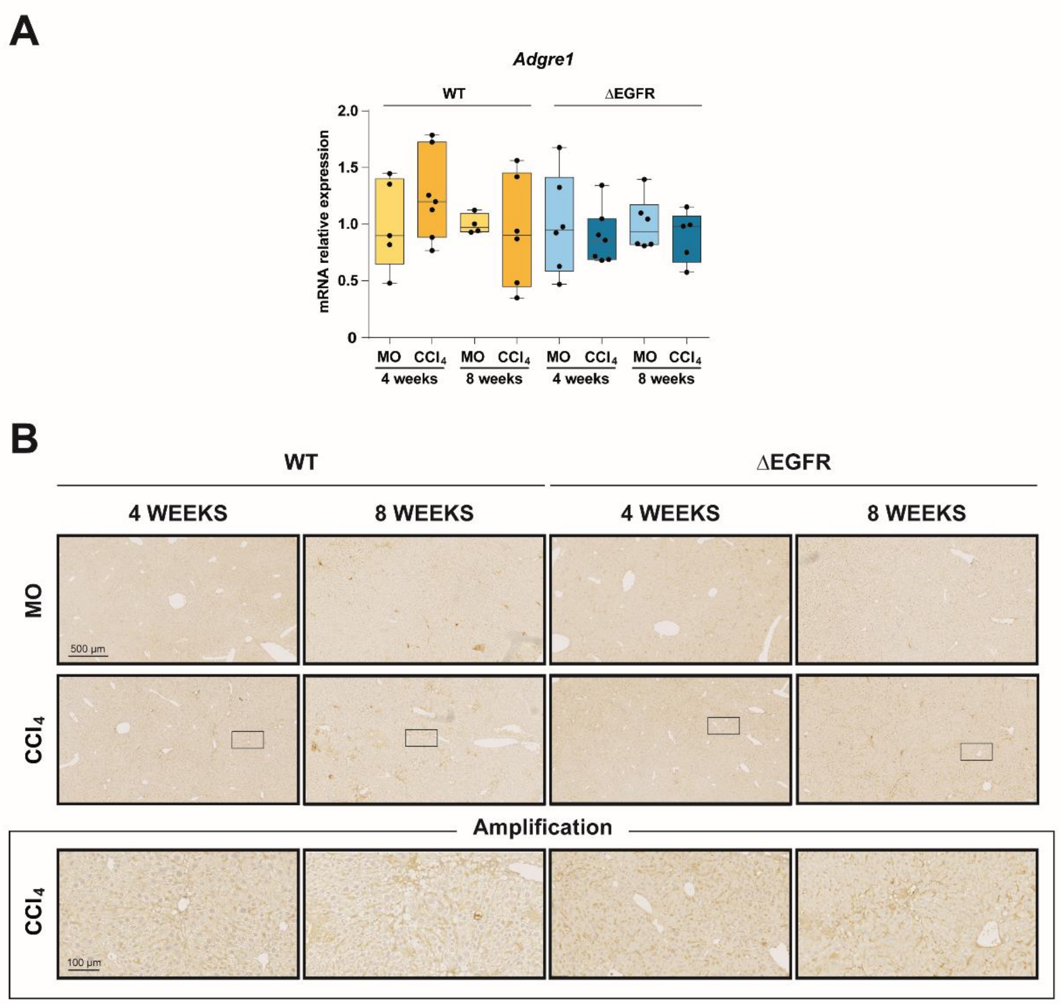


**Supplementary Figure 3. Macrophage infiltrate in livers from $CCl_4$-treated WT and ΔEGFR mice.** ΔEGFR and WT mice were treated with $CCl_4$ or vehicle (MO) for 4 or 8 weeks**. A.** The mRNA expression of *Adgre1* was assessed by RT-qPCR. **B.** F4/80 immunostaining of macrophages. Data (n=4-7 animals per group) were analyzed using Student's t test and presented as box-and-whisker plots. Scale Bars: 500 µm and 100 µm (amplification).

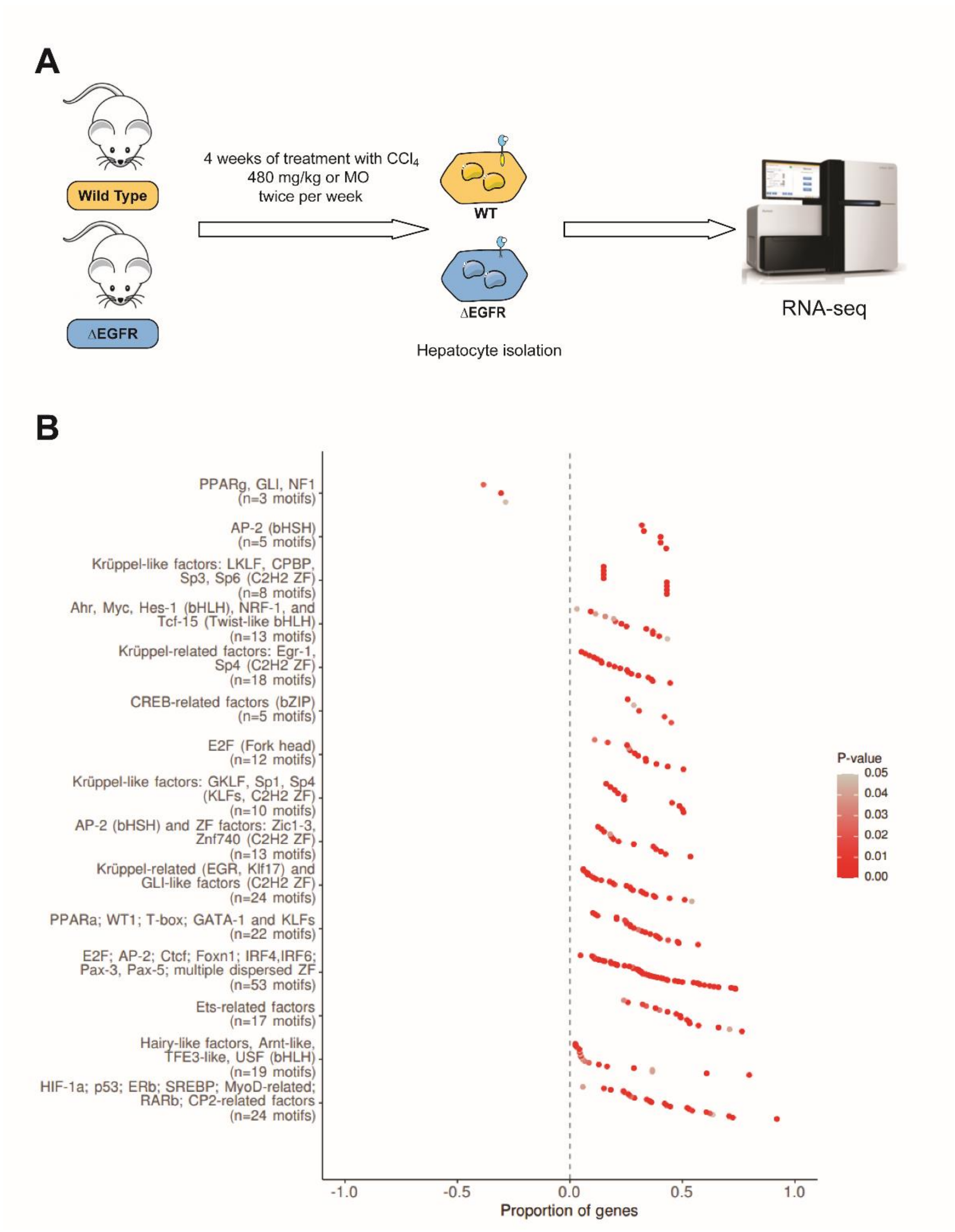


**Supplementary Figure 4. The EGFR pathway regulates the hepatocyte gene transcriptome during the response to $CCl_4$. Complementary information to Fig. 5. A.** Schematic representation of the workflow followed for the RNA-seq analysis conducted in this manuscript. Hepatocytes were isolated from ΔEGFR and WT mice treated with $CCl_4$ or vehicle (MO) for 4 weeks (4 animals per group, excepting WT-$CCl_4$, which were 3). **B.** Analysis of enriched transcription factors (TF) activity. Proportion of genes shows the ratio between the set of differentially expressed genes with the total size of genes in the gene set/motif. Dots on the left side of the plot show TF enriched in ΔEGFR condition, while dots on the right side show TF enriched in WT condition. Significant TF motifs are clustered based on common over-represented genes. Data were analyzed using an over-representation analysis against TRANSFAC database.

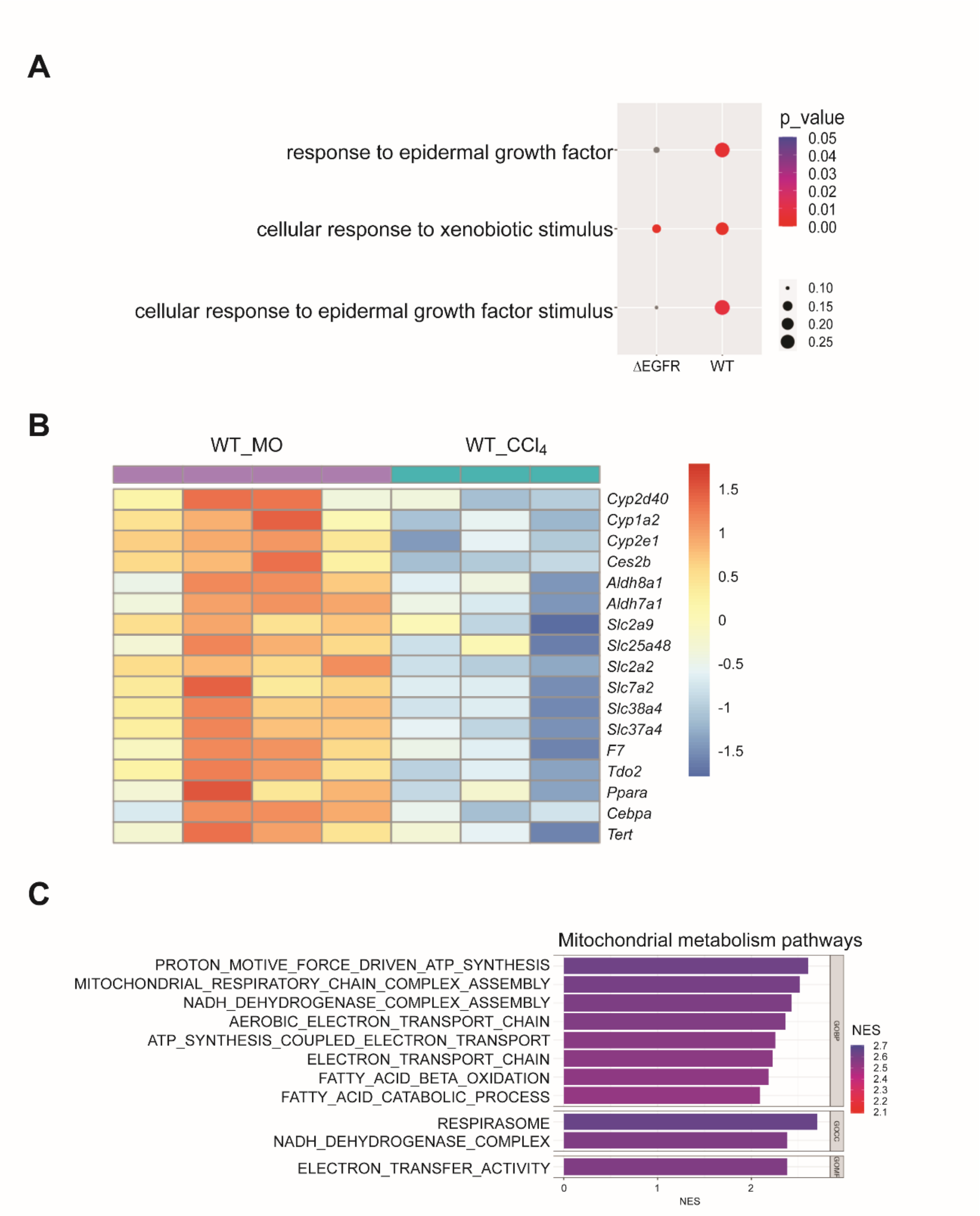


**Supplementary Figure 5 The EGFR pathway regulates the hepatocyte gene transcriptome during the response to $CCl_4$. Complementary information to Fig. 5. A.** Dot plot showing differences in enrichment for key events related to xenobiotic and epidermal growth factor stimulus between ΔEGFR and WT mice. **B.** Heatmap showing changes in the expression of genes related to liver-specific functions that appeared specifically in WT, but not in ΔEGFR, hepatocytes after $CCl_4$ treatment. **C.** Barplot of mitochondrial metabolism pathways that appeared enriched when comparing treated hepatocytes, ΔEGFR versus WT, by GSEA. Data (4 animals per group, excepting WT-$CCl_4$, which were 3) were analyzed using pre-ranked GSEA.

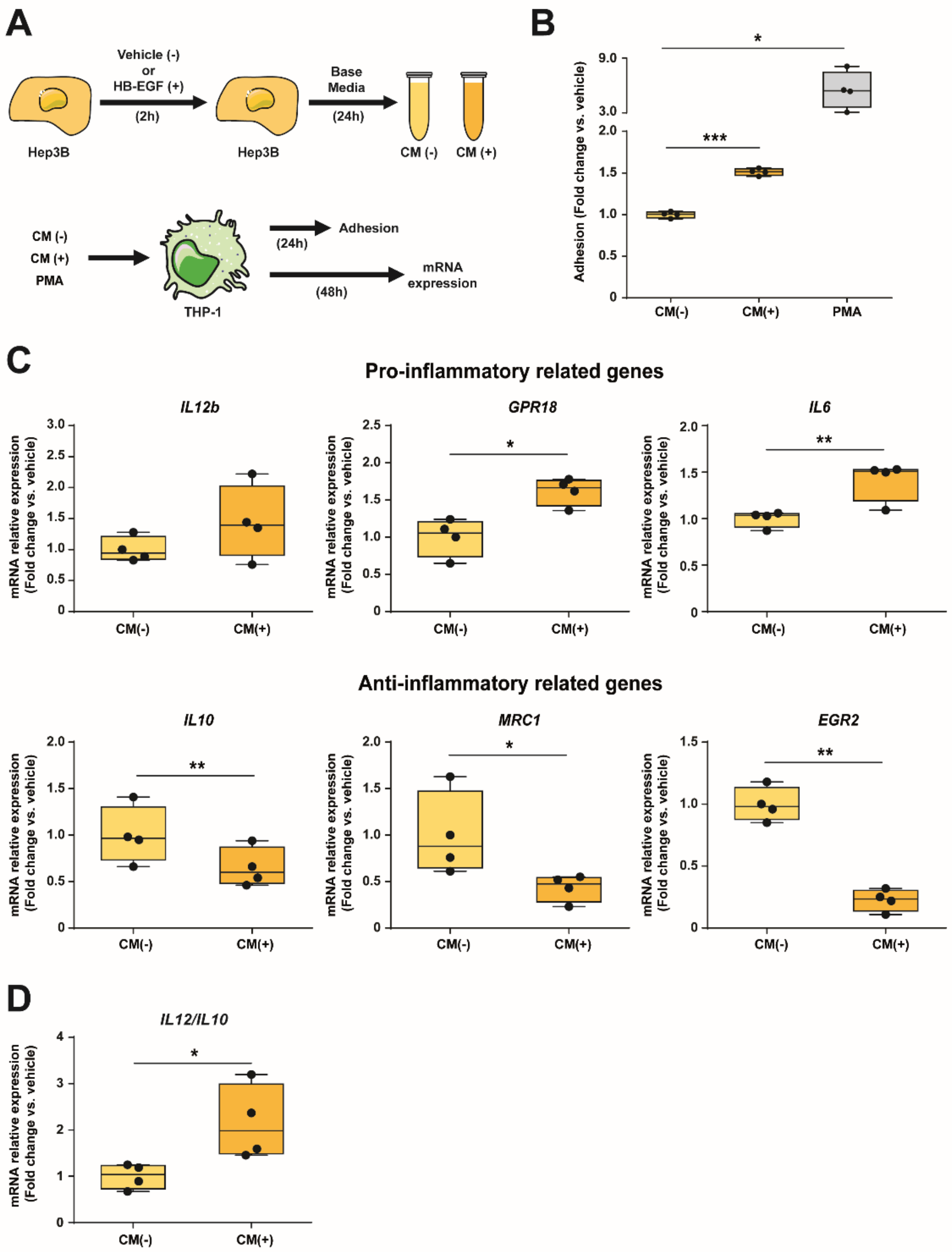


**Supplementary Figure 6. The EGFR pathway regulates the human hepatocyte secretome. Complementary information to Fig. 6.** Hep3B cells were used as a human hepatic model and THP-1 as a human monocyte cell line. **A**. Schematic representation of the experimental approach followed to collect conditioned media and analyze their effect on human macrophages phenotype. **B**. Effect of conditioned media from Hep3B cells treated with (CM+) or without (CM-) HB-EGF (20 ng/ml) on the adhesion of human monocyte cell line THP-1. phorbol myristate acetate (PMA) was used as positive control. **C.** The expression levels of M1 and M2 associated genes in the THP-1 cells were measured by RT-qPCR. **D.** Il12/Il10 ratio was used as indicator of the pro/anti-inflammatory phenotype of macrophages. Data (from at least three independent experiments) are expressed as fold change versus vehicle (Conditioned media from untreated Hep3B cells, CM-) and presented as box-and- whisker plots. *p<0.05, **p<0.01, ***p<0.001 using Student's t test.
