## Supplementary figures and images for "The hepatocyte Epidermal Growth Factor Receptor (EGFR) pathway regulates the cellular interactome within the liver fibrotic niche"

### Visual Abstract

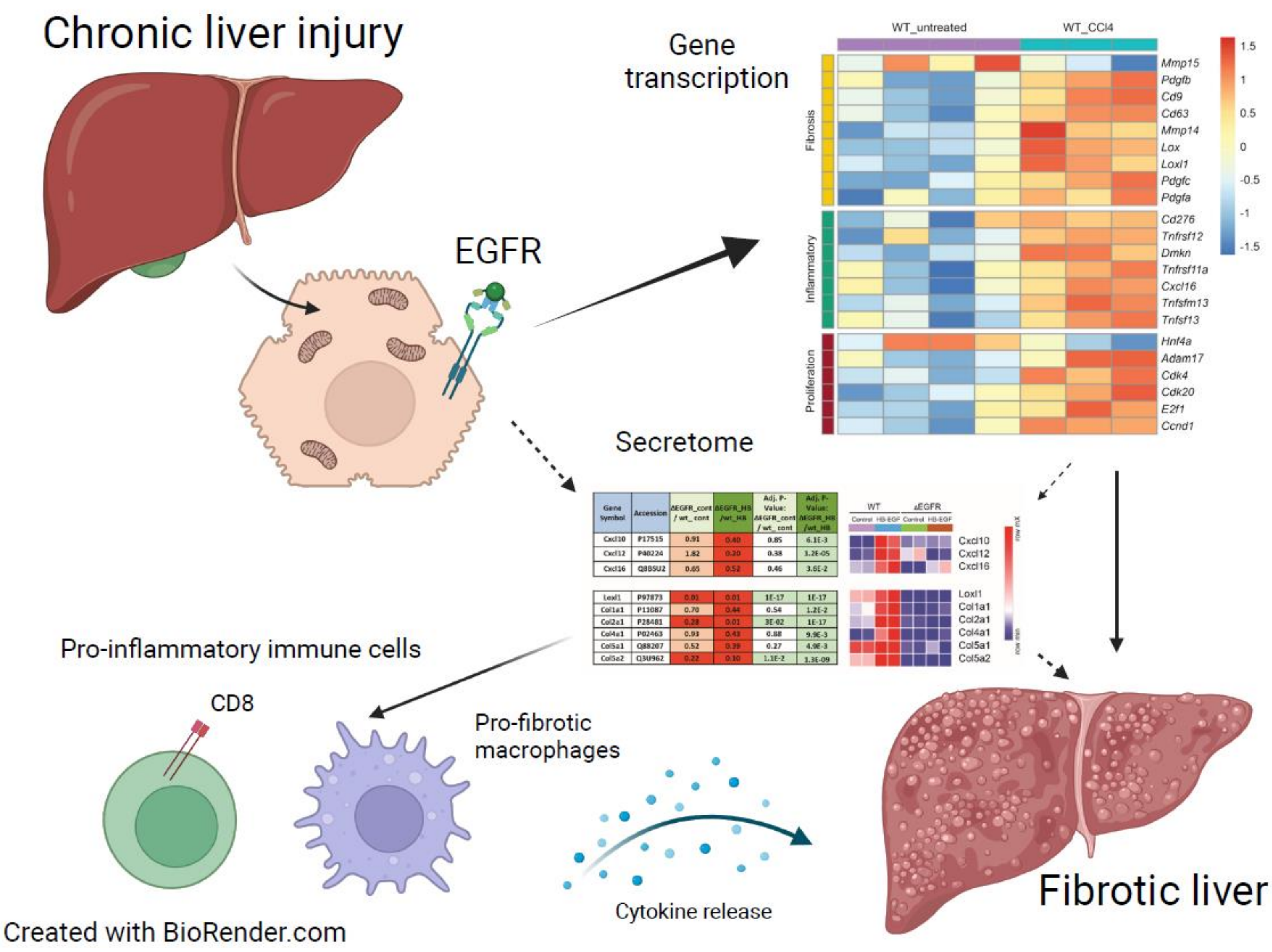


Gonzalez-Sanchez, Vaquero et al.
